# A Sequential Triple-Drug Strategy for Selective Targeting of p53-Mutant Cancers

**DOI:** 10.1101/2025.11.06.687028

**Authors:** Mohammed M. Alruwaili, Yanqi Guo, Justin Zonneville, Thomas Melendy, Robert M. Straubinger, Barbara A. Foster, Priyanka Rajan, Henry Withers, Sarah Chatley, Renuka Iyer, Christos Fountzilas, Andrei V. Bakin

## Abstract

The tumor suppressor *TP53* gene (p53) is mutated in most human malignancies; however, existing treatment options are largely ineffective, lack selectivity, and cause toxic side effects. To address these clinical problems, we developed a sequential triple-drug strategy for p53 mutant cancer cells. Here we show that a combination of a thymidine analogue (TAS102) plus PARP inhibitor (PARPi) promotes formation of DNA double-strand breaks (DSBs) and G2-arrest specifically in p53 mutant cancer cells. Transcriptome analysis revealed that TAS102-PARPi treatment of p53 mutant cells did not repress DNA replication but activated DSB repair and blocked the mitotic program, consistent with G2-arrest. In contrast, TAS102-PARPi treatment of normal p53 wild-type cells resulted in a temporal G1-arrest and rapid recovery of cell cycle capacity after drug withdrawal. In p53 mutant cancer cells, subsequent blocking of a G2-checkpoint kinase, such as WEE1, released these G2-arrested cells into mitosis, leading to massive cell death. Delayed administration of a G2-kinase inhibitor provides time for p53 wild-type cells to repair DNA, thereby minimizing toxicity to normal tissues. This sequential triple-drug strategy exhibited robust efficacy in preclinical models of colorectal and pancreatic cancers and was well tolerated in mice. Together, our findings illustrate a promising triple-drug strategy for targeting p53 mutant malignancies.

## INTRODUCTION

Colorectal cancer (CRC) and pancreatic ductal adenocarcinoma (PDAC) are among the leading causes of cancer-related mortality worldwide, posing significant global health challenges^1^. Despite improvements in early detection ^2^, CRC remains a considerable clinical burden, as the majority of patients are diagnosed at advanced stages or progress to metastatic disease after initial treatment^3^. The 5-year overall survival (OS) rate for patients with metastatic disease is less than 15% ^4, 5^. PDAC is an aggressive cancer characterized by rapid disease progression, early metastasis, and limited response to conventional therapies^6^. The 5-year OS rate for PDAC for all stages combined remains under 10% despite the use of multi-agent chemotherapy regimens ^6, 7^. For patients with chemotherapy-refractory CRC or PDAC, there are currently no FDA-approved therapies that extend overall survival beyond 3 months^5, 7^. These poor clinical outcomes suggest the need for more effective therapeutic strategies that target the molecular vulnerabilities of these cancers.

The tumor suppressor *TP53* gene is mutated in most cancers, including CRC and PDAC, and mutant p53 (p53mut) contributes to cancer progression and poor response to therapy^8, 9^. p53 protein is critical for maintaining genomic stability by regulating DNA repair and cell cycle control at the G1/S checkpoint in response to DNA damage^10^. Despite ample knowledge of the functional consequences of p53 mutations, therapeutic approaches targeting p53mut cancers have been largely unsuccessful, and p53 status is largely ignored in the clinical management of patients^11^. Cancer cells with mutant p53 show greater reliance on the G2 checkpoint due to the loss of G1-checkpoint control. This vulnerability of p53mut tumors has been explored with agents targeting the G2 checkpoint alone or in combination with chemotherapeutics, such as doxorubicin and carboplatin^12–14^. The major limitations of these approaches are the low selectivity of these therapeutics for p53 status and the high hematologic and organ toxicities in combination therapy^15, 16^. Therapeutic strategies with enhanced selectivity towards p53mut cancers are needed to improve treatments of these malignancies ^10^.

Recently, we developed a two-drug therapeutic strategy that selectively induced DNA damage in p53mut cancer cells ^17, 18^. This strategy combines the thymidine analog TAS102, which acts as an inducer of DNA single-strand breaks (SSBs) through post-replicative DNA repair, and PARP inhibitor (PARPi) which acts as an amplifier of DNA damage by converting SSBs to more lethal double-strand DNA breaks (DSBs)^17, 18^. TAS102-PARPi treatment induces G1-arrest in wild-type p53 (p53WT) cells, while p53mut cells exhibit high DSB levels, G2-arrest, and increased cell death ^18^. The inducer-amplifier strategy showed greater efficacy in p53mut cancer models, including CRC and PDAC, compared to either drug alone, while no major toxicities were observed in mice ^17, 18^. This two-drug strategy is being tested in a first-in-human Phase I clinical trial for patients with refractory CRC using the TAS102-talazoparib combination (NCT04511039)^19^. Preliminary data showed no significant signs of toxicity and improved median progression-free survival (mPFS) compared to historical mPFS for TAS102 alone^19^.

Here, we show that TAS102-PARPi treatment induces prolonged G2-arrest in p53mut cancer cells even after drug washout, while non-tumor p53WT cells rapidly repair their DNA and re-enter the cell cycle. The ATR-WEE1 signaling axis is critical for G2-arrest in response to TAS102-PARPi in p53mut cancer cells. Following TAS102-PARPi treatment, a selective inhibitor of WEE1 kinase released p53mut cells with unrepaired DNA into mitosis, leading to cell death. This sequential approach combining TAS102-PARPi treatment with a WEE1 inhibitor showed greater tumor control in preclinical models than either treatment alone. The sequential triple-drug regimen was well tolerated in animals even after extended administration.

## MATERIALS AND METHODS

### Cell culture models

The human CRC cell lines HT29 (RRID:CVCL_0320) and DLD1 (RRID:CVCL_0248), human non-tumor breast epithelial MCF10A (RRID: CVCL_0598) and human breast cancer MDA-MB-231 (RRID:CVCL_0062) were obtained from the American Type Culture Collection (ATCC, Manassas, VA, USA) and cultured as recommended by ATCC. The human CRC cell lines HCT116, HCT116-p53(-/-; KO), HCT116-p53(R248W/-), SW48, and SW48-p53(-/-; KO) were obtained from Dr. Bert Vogelstein, Johns Hopkins University. Human CRC RKO and RKO-p53(-/-, KO) cell lines were obtained from Dr. Alessandro Carugo, IRBM European School of Molecular Medicine, Italy. The details of the genetic information for the cell line and patient-derived xenograft (PDX) models are presented in **Key Resources Table**. All human cell lines were authenticated using short tandem repeat profiling by ATCC or the Roswell Park Genomic Shared Resource within the last three years. The cells were routinely screened for mycoplasma, and all studies made use of mycoplasma-free cells. Cell cultures were maintained in media and incubated as recommended by ATCC.

### Cell-derived xenografts in mice

Female SCID/CB17 mice (6−8-week-old) were obtained from a colony of SCID/CB17 mice that were bred and maintained at the Animal Facility of the Roswell Park Comprehensive Cancer Center. Animals were kept in microinsulator units and were provided with food and water *ad libitum* according to a protocol and guidelines approved by the Institute Animal Care and Use Committee (IACUC). The facility was certified by the American Association for Accreditation of Laboratory Animal Care (AAALAC) and in accordance with the current regulations and standards of the US Department of Agriculture and the US Department of Health and Human Services. Mice were inoculated subcutaneously into the left flank with exponentially growing HT29 tumor cells at (1×10^6^/mouse). Tumor growth was monitored by measuring tumor diameters with electronic calipers twice/a week. The volume was calculated using the formula (length) × (width)^2^/2. The weights of mice were measured twice per week. Once the tumor volume reached 50-100 mm^3^, mice were randomly assigned (n = 8-10/group) into four treatment arms: (1) vehicle control, (2) WEE1 inhibitor (WEE1i, adavosertib, MK-1775, 40-60 mg/kg), (3) TAS102-talazoparib (TAS102: 50 mg/kg, talazoparib: 0.150 mg/kg), and (4) sequential regimen (TAS102-talazoparib for one week followed by WEE1i in the second week). TAS102 and talazoparib were dissolved in 12% HPCD (2-hydroxypropyl)-β-cyclodextrin in DPBS, whereas MK-1775 was prepared in 5% DMSO, 40% PEG300, 5% Tween-80, and 50% saline. All drugs were freshly prepared and administered via oral gavage on a 5-days-on, 2-days-off schedule. At a tumor diameter of 2cm^3^, mice were euthanized and subjected to necropsy and organ collection. Tumor growth assessment was stopped once a mouse reached the endpoint (typically in a vehicle-treated cohort) while survival data were still collected in the study. Tumor tissues were collected by snap-freezing in liquid nitrogen for RNA and protein analyses. Blood was collected for CBC by cardiac puncture.

### Patient Derived Xenograft (PDX)

Donor and experimental pancreatic patient-derived xenografts (PDAC-PDX-14312-4p) were implanted subcutaneously into SCID mice, whereas colorectal patient-derived xenografts (CRC PDX-3B1-29-RP-3p; APC E1353*, KRAS-G12D, TP53-R213*, PIK3CA-E545K) were grown in NOD-Cg-Prkdcscid Il2rgtm1Wjl/SzJ mice (Jackson Laboratory, Bar Harbor, ME, USA), also known as NOD SCID gamma (NSG). Tumors were grown in donor mice to a volume of ∼1500mm^3^, harvested rapidly after euthanasia, cut into 2×2×2 mm^3^ pieces under sterile conditions, and surgically implanted into the left flank. Surgical staples were removed about ten days post implantation, and tumor growth and mouse weight were measured twice per week. At the average tumor volume of 100mm^3^, mice were randomized and treated as described above.

### Complete Blood Counts

Blood was collected by cardiac puncture into a tube containing EDTA to prevent coagulation at the endpoint of the study. The analysis was performed using the HemaTrue Analyzer and HeskaView Integrated Software version 2.5.2. Graphs for the blood compounds were generated using GraphPad Prism8 (Version 8.4.2).

### Immunoblotting

Whole-cell lysates were prepared using NP40 Lysis Buffer (0.88% NP-40, 132 mM NaCl, 44 mM HEPES, and 8.8 mM NaF) supplemented with 2 mM sodium orthovanadate, 1 mM PMSF, and Protease Inhibitor Cocktail. Protein concentrations were quantified using the Bio-Rad DC Protein Assay (Cat# 5000111). Equal amounts of protein from each sample were heated for 5 min and resolved by SDS-PAGE at a constant voltage of 100V. Proteins were then transferred onto a nitrocellulose membrane (0.2-µm pore size) at a constant voltage of 100V for 120 min at 4°C. Membranes were blocked in 1x TBS-T (Tris-buffered saline (TBS)) with 0.1% v/v TWEEN-20) and 5% w/v non-fat dry milk. Primary antibodies were diluted in 5% BSA or 5% milk in 1x TBS-T and incubated overnight at 4°C. The membranes were washed three times with 1x TBST for 10 min each and then incubated with secondary mouse or rabbit HRP-conjugated antibody for 1h at room temperature. The membranes were washed three times with 1x TBST for 15 min each. Pierce ECL Western Blotting Substrate was used for chemiluminescent detection. Signals were visualized and imaged using radiographic films.

### Cell cycle analysis by Flow Cytometry

Cells were seeded at 300,000 cells/well in 6-well plates and then treated for 24h with various concentrations of drugs, listed in **Key Resources Table**. Cells were collected and fixed with ice-cold 70% ethanol for 1h at -20°C and stained for 1h at 4°C in Krishan DNA Buffer (propidium iodide, sodium citrate, RNase A, NP40, and 0.1 mM HCl). The samples were analyzed on a BD LSR Fortessa cytometer running FACS Diva (version 6.1.3), and the data were processed using FCS Express 7 (version 7.06.0015). The experiments were repeated twice, and representative histograms are shown.

### Cytotoxicity

The cells were seeded at a density of 5,000 cells/well in a 96-well plate. After 24h, cells were treated with TAS102 at a range of concentrations in the presence of 100nM talazoparib, and/or the G2 kinase inhibitors at various concentrations for 72h. The cells were stained with 1% methylene blue for 30 min, rinsed with water, and dried overnight. The cells were solubilized in 5% SDS prepared in PBS, and absorbance was measured at 650 nm. IC50 values were calculated using GraphPad Prism (version 10.0.3). Combination Index (CI) values were calculated based on the Loewe Additivity Model as implemented in the SynergyFinder R package using the following formula: CI = D1/IC50_1_ + D2/IC50_2_, where D1 and D2 are the concentrations of drugs 1 and 2 used in the combination, while IC50_1_ and IC50_2_ are the concentrations of drugs 1 and 2 inhibiting 50% of the activity when used individually. Each dot in the Scatter plot represents the drug interaction in the combination at specific concentrations, plotted against the fraction affected (Fa). The CI values are distributed below, at, or above the threshold of 1, indicating synergy (CI < 1), additivity (CI = 1), or antagonism (CI > 1).

### Immunohistochemistry (IHC)

Freshly isolated tumors were fixed in 10% neutral buffered formalin (Sigma-Aldrich, Cat. #HT501128) for 48h, then transferred to 70% ethanol for dehydration, before embedding in paraffin at Roswell Park’s Pathology Network Shared Resources (PNSR) into formalin-fixed paraffin-embedded (FFPE) blocks. Staining for hematoxylin & eosin (H&E) and for p-Y15-CDK1, Cyclin B1, and p-S10-HH3 was performed by the PNSR. Antibodies and reagents are listed in **Key Resources Table.** Slides were scanned by Aperio ImageScope version 12.4.0.5043, and representative images were captured at 20x magnification. Three representative tumors from each treatment group were used for quantification. A minimum of eight fields of view (FOV) were analyzed for each tumor sample. Entire stained tissue sections were scanned, and the areas of highest density for immunoreactivity (“hot spots”) were manually selected for quantification. Each “hot spot” was recorded at 20x magnification. The positive area was calculated using the formula: Negative area – Positive area × 100) in ImageJ. Data were reported as mean and standard error (mean +-SE). Data are presented using GraphPad Prism8 (v8.4.2).

### Annexin-V staining

Following drug exposure, the cells were trypsinized, washed with PBS, and resuspended in annexin-binding buffer. Dead cells were stained for annexin-V and DNA with propidium iodide according to the manufacturer’s instructions (Dead Cell Apoptosis Kit with Annexin-V-FITC and PI, for flow cytometry, ThermoFisher Scientific, Cat. No.: V13242). The samples were processed on a BD LSR Fortessa cytometer running FACS Diva (v6.1.3), and the data were analyzed using FCS Express 7 (v7.06.0015). Experiments were repeated twice, and representative histograms were shown.

### Immunofluorescence microscopy

The cells were grown on glass coverslips for 24h and then treated with 500nM TAS102, 100nM talazoparib, or their combination for the indicated time; alternatively, cell were treated with TAS102-talazoparib for 25h following drug withdrawal in incubation for indicated time. Cells were fixed with 4% PFA and permeabilized with 0.05% Triton X-100. Samples were blocked with 3% milk in PBS and probed with γH2AX and RAD51 antibodies in 1% milk, followed by a wash in PBS and incubation with fluorescent secondary antibodies in 0.5% milk. Samples were washed with PBS three times and DNA was labeled with Hoechst 33342 before mounting on glass slides. Fluorescence images were taken with a Plan Apochromat 100x objective (1.40 NA mineral oil) at ambient temperature using a Nikon TE2000-E inverted microscope equipped with a charge-coupled device camera (CoolSNAP HQ; Photometrics). The levels of γH2AX and RAD51 were assessed by measuring cellular fluorescence intensity as the Corrected Total Cellular Fluorescence (CTCF). Briefly, a minimum of six random fields per treatment group were evaluated using MetaVue imaging software (version 7.7.3, Molecular Devices). Three background readings were measured for each field of view and individual cells that were positively stained with γH2AX and RAD51 were selected with a region freehand tool. For each treatment group, >40 cells were evaluated and the CTCF values were calculated using the formula: integrated density – (area of cell × average background fluorescence). The CTCF values are reported as mean and standard error (mean +-SE). Data are presented using GraphPad Prism8 (v8.4.2).

### RNA sequence

Cells were seeded at a density of 1x10^6^ cells in a 100mm dish, and the following day media was replaced with base media containing 5% FBS for 6h. Cells were then treated with 500nM TAS102, 100nM talazoparib or their combination for 24h. Next day, cells were lysed with TRIzol reagent (Thermofisher Scientific; Cat # 15596026). RNA was isolated using Direct-zol RNA Miniprep Kit (Zymo Research; Cat # R2050). The sequencing libraries are prepared from 500ng total RNA using the mRNA HyperPrep kit (Roche) and following manufacturer’s instructions. The polyA+ RNA population was captured using the mRNA Capture Beads and then utilized for cDNA synthesis using random primers and reverse transcriptase. cDNA was converted into blunt-ended double-strand DNA using Taq pol (overhanging A) and purified using Pure Beads (Roche). Multiple indexing adapters with overhanging T at the 3’ end were ligated to the ds-cDNA ends. Adapter ligated ds-cDNA libraries were amplified by PCR, purified using Pure Beads, and then were validated for appropriate size on a 4200 TapeStation D1000 Screentape (Agilent Technologies, Inc.). The DNA libraries were quantitated using KAPA Biosystems qPCR kit, and were pooled together in an equimolar ratio, following experimental design criteria. Each pool was denatured and diluted to 200pM with 1% PhiX control library added. The resulting pool was then loaded into the appropriate NovaSeq Reagent cartridge, as determined by the desired sequencing depth and number of samples. 100-cycle paired-end sequencing was then performed on a NovaSeq6000 following the manufacturer’s recommended protocol (Illumina Inc.).

Raw count files were generated from FASTQ files using UB CCR’s HPC nr-core/rnaseq pipeline (https://nf-co.re/rnaseq/3.12.0). Raw sequencing reads passed quality filters from Illumina RTA were first pre-processed by using FASTQC (v0.11.8) for sequencing base quality control. The reads were mapped to GRCh38 human reference genome and the GENCODE (v38) (128) annotation database using STAR (v2.7.9a). Counts from RSEM were filtered and then upper quartile normalized using R package edgeR. Gene Ontology analysis of biological processes pathways was selected based on adjusted p-values, and the NES was calculated by ranking all genes based on their differential expression, calculating the Enrichment Score (ES) for each gene set, and normalizing the ES by dividing it by the mean ES values from random permutations. Cell-cycle genes in the heatmap were derived from the Cyclebase 3.0 database.

### Quantification and Statistical Analysis

Statistical analysis was performed using GraphPad Prism software (v10.0.3) unless otherwise indicated. The data represent biological replicates (n) as mean value with standard error (mean ± SE). Statistical significance of data comparisons was determined using the Student’s unpaired *t*-test with a two-tailed distribution, one way ANOVA, or two-way ANOVA with a Tukey multiple comparisons test or with Sidak’s multiple comparisons test as indicated in figure legends. Statistical significance was achieved when *P*<0.05. The correlation analysis was performed using the Pearson correlation approach in RStudio. Survival was evaluated using the Kaplan-Meier estimator with the log-rank test, based on time-to-arrive at a defined tumor volume using GraphPad Prism. Cytotoxicity data were normalized to that of the control, and IC50 values were calculated using three-parameter nonlinear regression.

## RESULTS

### The TAS102-talazoparib treatment activates DNA damage response and arrests cells in G2

Prior studies have shown that the incorporation into DNA of thymidine analogs (i.e., ethynyl-deoxyuridine, EdU, or trifluorothymidine, a component of TAS102) triggers post-replicative repair, resulting in single-strand DNA break (SSB) intermediates ^17, 18^. The addition of a PARP inhibitor (PARPi) greatly increases the formation of double-strand breaks (DSBs) ^17, 18^. DNA damage activates the p53-dependent G1 checkpoint and DNA repair in non-tumor p53WT cells, whereas p53mut cells (exhibiting a compromised G1 checkpoint) accumulate DSBs and respond with the G2 checkpoint ^17, 18^. To define the molecular mechanisms of the G2 checkpoint response to TAS102-PARPi treatment, isogenic CRC cell lines with p53WT, p53mut or p53-deletion (KO, knockout) were utilized. Treatment with talazoparib (PARPi) for 24h increased the G2 population in p53mut and p53KO cells but had little effect on p53WT cells, whereas TAS102 alone increased the G2 population in p53mut/p53KO cells and a moderate G2 increase in p53WT cells (**Fig. 1a-b**). TAS102-talazoparib treatment (TAS-Tal) exerted in p53mut/p53KO cells a strong increase in the G2 population and a reciprocal decrease in the G1 population, and these changes were statistically significant (**Fig.1a-b**). In p53WT cells, the combination did not increase the G2 population compared to TAS102 alone. Immunoblot analysis of cell cycle and DNA damage response (DDR) markers showed that TAS102 and TAS102-PARPi decreased the activating phosphorylation of PLK1, a kinase mediating mitotic transition, independent of p53 status (**Fig.1c-d**). TAS102-PARPi treatment increased an inactivating phosphorylation of CDK1 at Tyr15 (**Fig. 1c-d**), a site known to be phosphorylated by WEE1 kinase. Protein levels of cyclins A2 and B1, partners of CDK1, were also elevated (**Fig.1c-d**). In contrast, phosphorylation of histone H3 was not increased, indicating no entry into mitosis (**Fig.1c-d**). Together, these results showed induction of the G2 checkpoint and DDR in p53-deficient cells upon treatment with TAS102 alone and more robustly in combination with PARPi (**Fig.1g**). To validate this observation *in vivo*, the G2 markers were examined by immunohistochemistry (IHC) in tumor tissues from the CRC PDX models (generated within the Phase I clinical NCT04511039 study), which were treated with vehicle or drugs (talazoparib, TAS102, or combination) as described ^18^. IHC analysis showed that TAS102 and TAS102-PARPi increased the levels of phospho-Tyr15-CDK1 and Cyclin B1 in p53^H179R^ mutant model (PDX-02-07-RP), whereas the levels of phospho-H3 were reduced (**Fig. 1e** and **Suppl. Fig.1a-b**). Similar results were obtained in the PDX-01-03-RP model carrying p53WT, although the effects of the drug combination were lower in the p53WT model (**Fig. 1f** and **Suppl. Fig.1c-d**). These findings indicate that the TAS102-PARPi combination primarily activated the G2 checkpoint in p53mut cancers, while p53WT cells responded by activating both G1 and G2 checkpoints (**Fig. 1g**).

**Figure 1.**
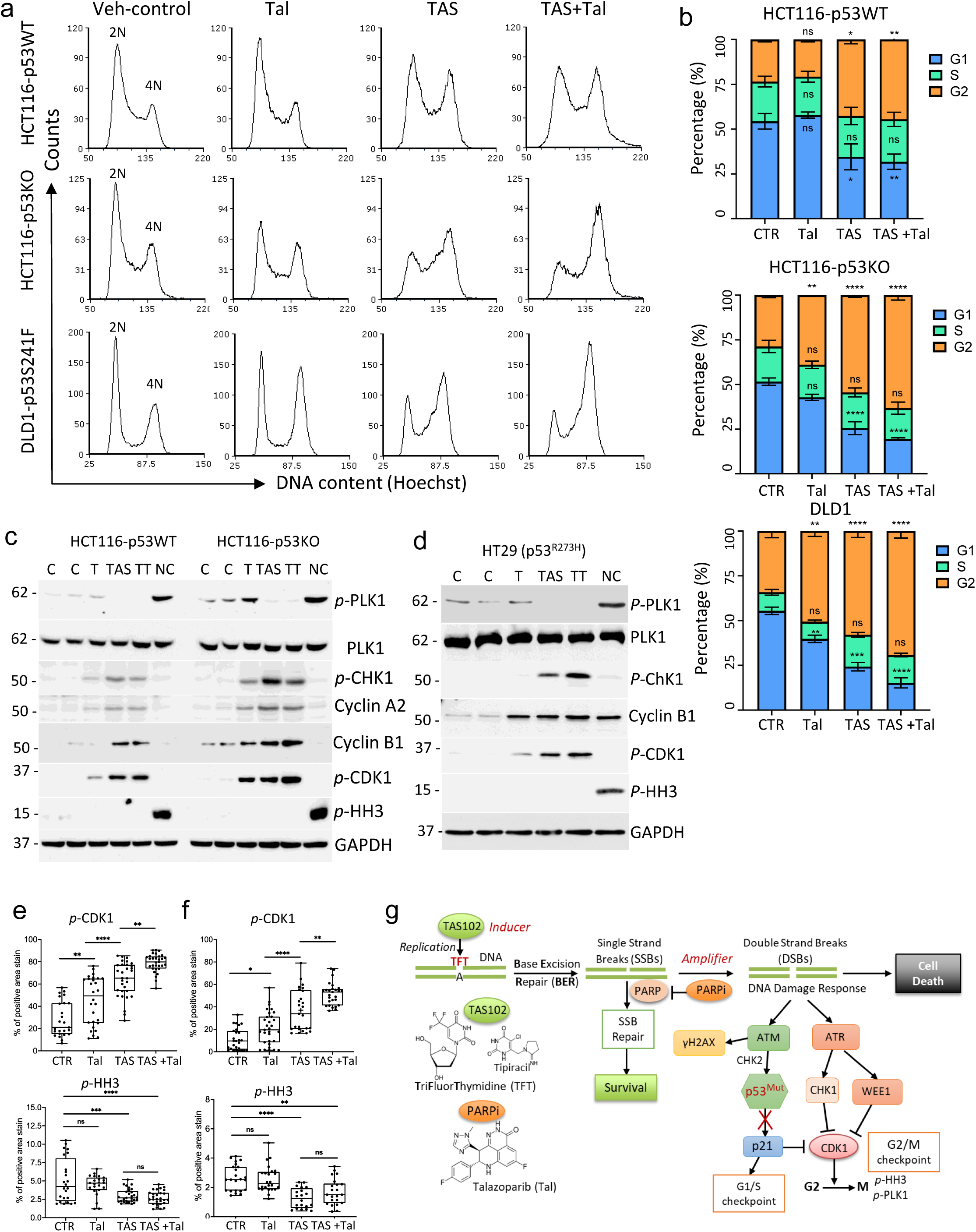
The combination of TAS102 and talazoparib induces DNA damage response and the G2 checkpoint. (**a-b**) Cell-cycle analysis of HCT116 p53WT and p53KO, and DLD1 (p53^S241F^) cell lines treated with vehicle control (C), 100nM talazoparib (Tal), 500 nM TAS102 (TAS) or their combination (TAS+Tal) for 24 hours. Cells were stained with Hoechst for DNA content and analyzed by flow cytometry. (**b**) Quantification of cell cycle data. Comparisons were made using two-way ANOVA (∗p < 0.05, ∗∗p < 0.01, ∗∗∗p < 0.001, ∗∗∗∗p < 0.0001). Experiments were repeated at least two times. (**c-d**) Immunoblots of whole-cell extracts from cell lines treated with vehicle control (C), 100 nM talazoparib (T), 500 nM TAS102 (TAS), their combination (TT), or 50 nM nocodazole (Nc) for 24 hours. (**e-f**) Quantification of phospho-CDK1 and phospho-H3 levels in CRC PDX-02-07-RP (p53^H179R^) tissues (**e**), and PDX-01-03-RP (p53WT) tissues (**f**). PDX tumor bearing mice were treated by oral gavage with vehicle-control, talazoparib, TAS102, or TAS102-talazoparib. Comparisons were done using one-way ANOVA (*p < 0.05, **p < 0.01, ***p < 0.001, and ****p < 0.0001). (**g**) Working model.

### TAS102-talazporib alters the transcriptomic program in p53WT and p53mut cells

To better define the response to the two-drug combination, transcriptomic analysis was performed on isogenic HCT116 p53WT and p53KO cell lines treated with drug monotherapies or their combination for 24h. Talazoparib alone induced minimal transcriptomic changes in both cell lines (**Fig. 2a, d**). In contrast, TAS102 exerted strong effects by inducing expression of 774 genes and downregulating expression of 367 genes in p53WT cells and 614 upregulated and 418 downregulated genes in p53KO cells relative to vehicle-treated controls (**Fig. 2b, e**). The drug combination in p53WT cells increased expression of 897 genes and decreased expression of 368 genes, while in p53KO cells expression of 698 genes was induced and that of 615 genes was decreased (**Fig. 2d, f**). The Venn diagrams demonstrated a notable overlap in altered gene expression between TAS102 monotherapy and the TAS102-PARPi combination, with 936 genes shared in p53WT and 770 shared in p53KO cells (**Fig. 2g-h**).

**Figure 2.**
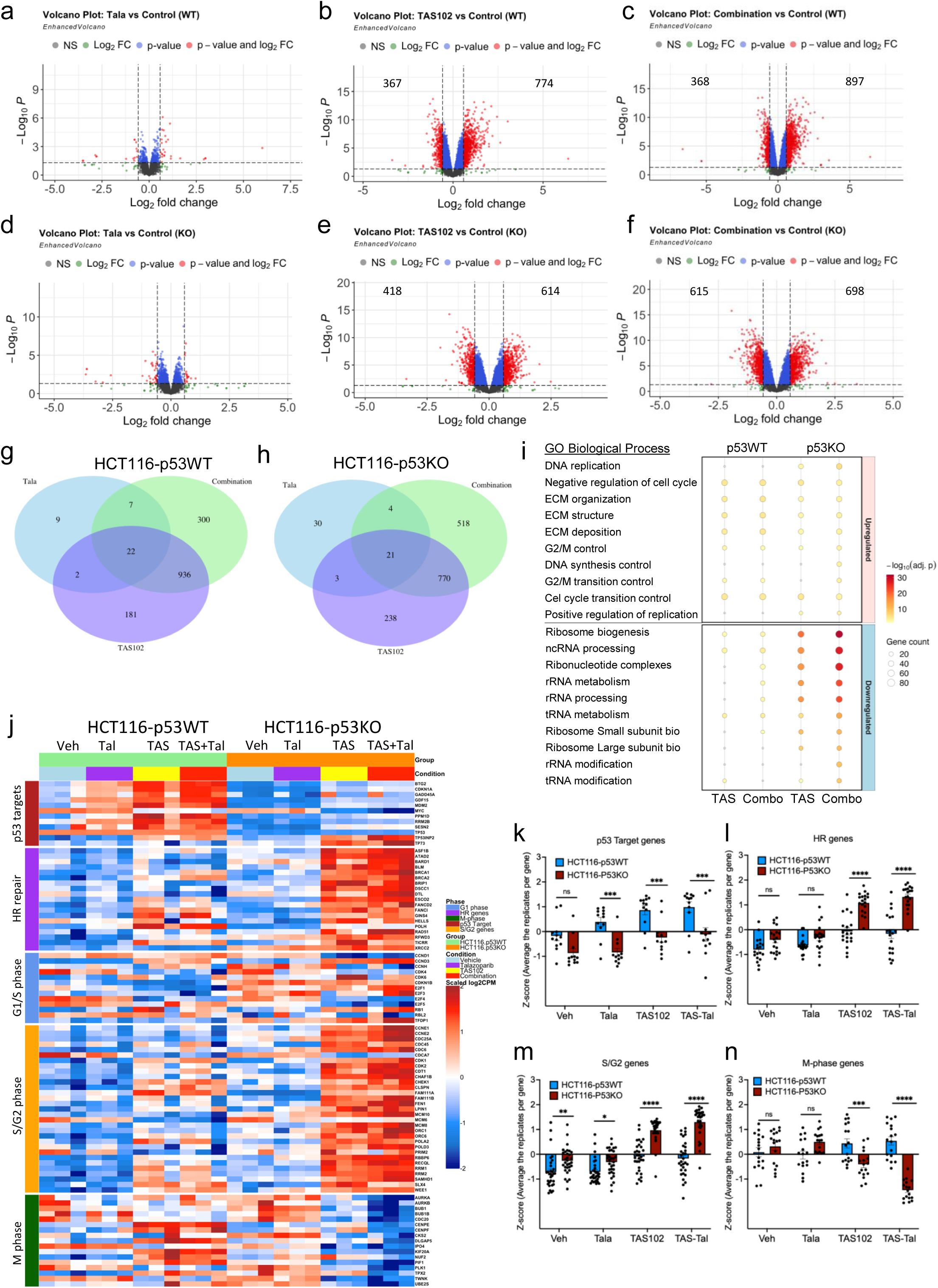
Transcriptomic changes in p53WT and p53KO cells following TAS102 and talazoparib treatment. Volcano plots show differential gene expression between control and treatment groups (talazoparib, TAS102, and combination) for both HCT116 p53WT (**a-c**) and p53KO (**d-f**) cell lines. Red data points represent genes that meet both the adjusted p-value <0.05 and a pre-defined log2 fold change (FC) threshold. Blue data points indicate genes that only meet the log2 FC threshold, gray dots show non-significant genes. (**g-h**) Venn diagrams show the overlap of differentially expressed genes among the treatment groups in HCT116 p53WT (**g**) and p53KO (**h**) cells. (**i**) Heatmaps show the expression profiles of select genes across the control and treatment groups in HCT116 p53WT (left) and p53KO (right) cells. Gene lists are curated from KEGG and Cyclebase_3.0 databases.

The gene ontology (GO) enrichment analysis was conducted on the differentially expressed genes (**DEGs**) to infer the biological processes affected by the drug treatments. The GO analysis showed that in both p53WT and p53KO cells treated with TAS102, the up-regulated genes were enriched in processes associated with cell cycle regulation and extracellular matrix (ECM) organization (**Suppl. Fig. 2a-b**). However, among the downregulated genes distinct processes were affected in p53WT *vs*. p53KO cells. In p53WT cells, downregulated genes were mainly associated with processes related to mRNA splicing and pyrimidine metabolism, whereas in p53KO cells with processes related to ribosome biogenesis, rRNA processing and maturation (**Suppl. Fig. 2a-b**). In TAS102-talazoparib treatment, the top upregulated biological processes in p53KO cells were related to DNA replication and G2/M transition, whereas downregulated processes were related to ribosome biogenesis (**Suppl. Fig. 2c-d**). The dot plots summarize the top significantly enriched processes related to cell cycle regulation, ECM organization, and ribosome biogenesis for p53WT and p53KO cells (**Fig. 2i**). The upregulation of processes related to ECM organization suggests a contribution of the matrix to cell division, while repression of ribosome biogenesis is consistent with reduced ribosome biogenesis at the G2 checkpoint in preparation for mitosis ^20^.

To reveal the transcriptome changes related to the cell cycle and DNA repair in p53WT *vs*. p53KO cells, heatmaps were built for p53 targets, cell cycle, and DNA repair genes identified in the GO analysis. The regulation of p53 target genes was clearly different between p53WT and p53KO cells (**Fig. 2j, k**). Treatment of p53WT cells with TAS102 (TAS) or TAS102-PARPi (TAS+Tal) strongly induced the expression of p53 targets (i.e., p21/CDKN1A, MDM2, BTG2); in contrast their regulation was markedly lower in p53KO cells (**Fig. 2j, k**, p53 Targets). In p53KO cells the expression of homologous recombination (HR) DNA repair genes (i.e., BRCA1, BRCA2, RAD51) was robustly induced by TAS102-PARPi treatment, whereas the effect was lower in TAS102 alone (**Fig. 2j, l**, HR genes). This finding is consistent with higher double-strand breaks (DSBs) induced by TAS102-PARPi treatment in p53KO cells compared to TAS102 alone ^17, 18^. The genes involved in G1/S transition (CDK4, E2F1, E2F3) and DNA replication (MCM6, PRIM2) were reduced in p53WT cells treated with TAS102 and TAS102-PARPi (**Fig. 2j, m**). In contrast, treatment of p53KO cells with TAS102 or TAS-PARPi increased the expression of genes mediating G1->S transition (E2F1, E2F3, TFDP1) and DNA replication (CCNE1, CDK2, MCM10, ORC1), **Fig. 2j, m**, G1/S and S/G2 genes. These effects show that neither TAS102 nor TAS-PARPi repress DNA replication in p53-deficient cells, whereas the treatments of p53WT cells induce a p53-dependent G1-arrest. Importantly, TAS-PARPi treatment of p53KO cells strongly reduced expression of genes involved in mitosis (AURKA, BUB1, CDC20, CENPF, PLK1, PIF1), an effect that was lower in TAS102 alone (**Fig. 2j**, **n**, M phase). In contrast, the expression of these genes was not reduced in p53WT cells. We further verified the transcriptome effects in the HT29 CRC cell line carrying p53^R273H^. RNAseq analysis showed that TAS102-PARPi treatment downregulates expression of M-phase genes and upregulates expression of HR-related genes comparable to the effects observed in HCT116-p53KO cells (**Suppl. Fig. 3**). The TAS102-PARPi effects were greater than those of TAS102 alone. Together, these data showed that the TAS102-PARPi treatment caused distinct transcriptome changes in p53-deficient *vs*. p53WT cells, consistent with G2-arrest and activation of HR-mediated DSB repair in p53mut cells and G1-arrest in p53WT cells.

### Targeting G2 checkpoint kinases and cytotoxicity of TAS102-PARPi treatment

We have shown that TAS102-PAPRi treatment activates signaling by ATM and ATR kinases ^18^, which are known to mediate DNA repair and G2-arrest ^21, 22^. Therefore, we examined whether blocking these kinases can enhance the toxicity of TAS102-PARPi in p53mut cells. A selective ATM inhibitor (KU55933; ATMi) did not enhance toxicity of TAS102-talazoparib in p53WT and p53-deficient (p53KO, p53^R248W/-^) HCT116 cell lines (**Suppl. Fig. 4a-c**). Instead, ATMi reduced sensitivity to TAS102-talazoparib in all tested lines, potentially due to ATM function in the G1/S checkpoint ^23^ and/or recovery of replication fork after DNA repair ^21, 22^. In contrast, the selective ATR inhibitor (AZD6738/ceralasertib; ATRi) markedly enhanced toxicity of TAS102-talazoparib in p53KO and p53mut cell lines (**Fig. 3a-d**). In p53-deficient cells the IC50 was enhanced 4-10-fold by 100nM ATRi (**Fig. 3b-d, n**), whereas only 1.5-fold increase was observed in p53WT cells (**Fig. 3a, n**). This result is consistent with our prior finding that, in cells treated with TAS102-talazoparib, blockade of ATR increases levels of γH2AX (a DNA damage marker) ^18^. Furthermore, blockade of ATR, but not ATM, increased phosphorylation of histone H3 (phospho-H3, a marker of mitosis) (**Suppl. Fig.4i**). These data are aligned with the role of ATR in resolving replication stress and activating the G2/M checkpoint, thus ensuring proper cell cycle progression and DNA repair ^24^. ATR kinase blockade allows cells with unrepaired DSBs to enter mitosis, leading to cell death ^22^.

**Figure 3.**
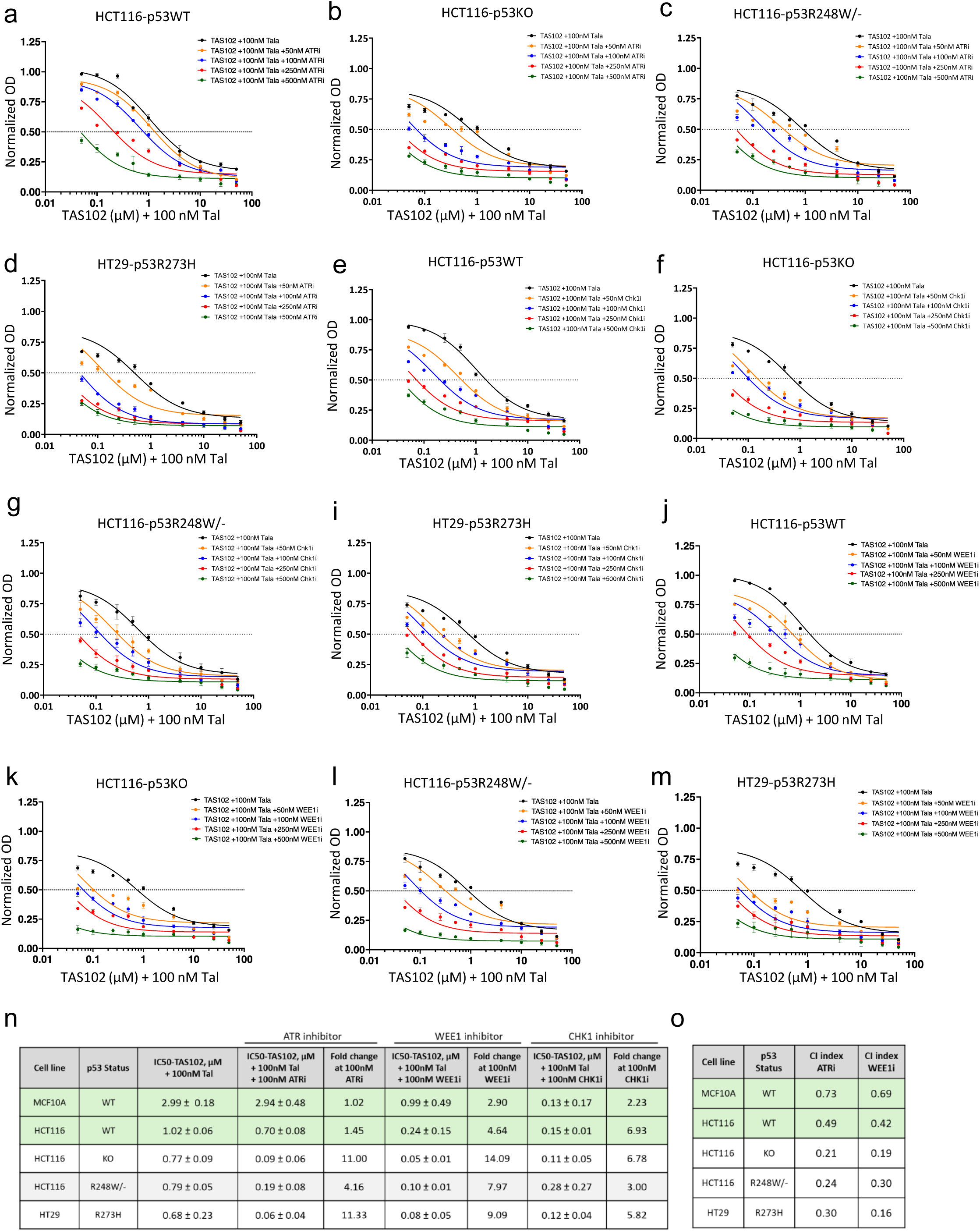
G2 checkpoint kinase inhibitors enhance the cytotoxic effects of TAS102 and talazoparib treatment. (**a-d**) Dose-response curves for a panel of colorectal cancer cell lines treated with increasing concentrations of TAS102 in the presence of 100 nM talazoparib. Cells were co-treated for 72 hours with varying concentrations of the ATR inhibitor AZD6738 (ATRi; 50nM, 100nM, 250nM, and 500nM). (**e-i**) Cells were treated as described above with the CHK1 inhibitor, AZD7762 (CHK1i; 50, 100, 250, and 500 nM). (**j-m**) Cells were treated as described above with the WEE1 inhibitor, MK-1775 (WEE1i; 50, 100, 250, and 500 nM). All data were normalized to untreated control. The half-maximal inhibitory concentration (IC50) values were determined using three-parameter nonlinear regression. All experiments were performed with at least two biological replicates and six technical replicates. (**n-o**) Cytotoxicity parameters in p53WT and mutant cell lines. Combination Index (CI) values were calculated for combinations with ATRi (**n**) and WEE1i (**o**) across several concentrations.

ATR kinase induces G2-checkpoint through downstream effector kinases such as CHK1 and WEE1; CHK1 kinase phosphorylates and inactivates CDC25 phosphatase, which controls CDK1, while WEE1 kinase inactivates CDK1 by phosphorylating Tyr15 ^21, 25–30^. Therefore, we explored whether blockade of these kinases would result in cytotoxic effects comparable to those of ATRi. A selective CHK1 inhibitor (AZD7762; CHK1i) showed comparable increase in cytotoxicity of TAS102-talazoparib in both p53WT and p53-deficient cells, about 8-10-fold at 100nM CHK1i (**Fig. 3e-i**). In contrast, a WEE1 inhibitor (MK1775; WEE1i) enhanced the cytotoxicity of TAS102-talazoparib preferentially in p53mut cells, 8-15-fold in p53mut *vs* 3-4-fold in p53WT cells at 100nM WEE1i (**Fig. 3j-m, o**).

Next, we examined the cytotoxicity of G2-kinase inhibitors in non-tumor p53WT MCF10A cells. ATRi and WEE1i showed only a moderate (2-3-fold) enhancement of toxicity of TAS102-talazoparib in non-tumor cells (**Suppl. Fig. 4d, f**). In contrast, CHK1i strongly enhanced toxicity of TAS102-talazoparib (nearly 10-fold) in non-tumor cells (**Suppl. Fig. 4e**). Immunoblotting verified the activities of CHK1 and WEE1 inhibitors on DNA damage response (DDR) markers in p53WT MCF10A and p53mut MDA-MB-231 cells (**Suppl. Fig. 4g, h**). Consistent with the effects of CHK1i *in vitro*, clinical trials have shown significant toxicities associated with CHK1i, resulting in discontinued early trials ^31, 32^. Conversely, drugs targeting ATR and WEE1 have shown tolerable toxicity in clinical trials ^33, 34^, positioning them as potential therapeutic candidates. Given that efficacy of TAS102-talazoparib was enhanced by both ATRi and WEE1i preferentially in p53mut cells, we evaluated the drug interaction using the Loewe Additivity Model ^35, 36^. The analysis revealed synergism between TAS102-talazoparib and both ATRi and WEE1i in p53mut cell models, with cooperativity index (CI) values ranging from 0.21 to 0.30 for ATRi (**Fig. 3n**, **Suppl. Fig. 5a-d**) and 0.16-0.30 for WEE1i (**Fig.3o**, **Suppl. Fig. 5f-i**). In contrast, in non-tumor p53WT cells, ATRi and WEE1i did not show substantial cooperativity with TAS102-talazoparib with the CI values of 0.73 and 0.69, respectively (**Fig. 3n-o** and **Suppl. Fig. 5e, j**). Thus, in this simultaneous drug regimen, TAS102-talazoparib synergized with ATRi and WEE1i in p53mut cancer cells, while exerting only a moderate interaction in p53WT cells.

### TAS102-talazoparib induces persistent DNA damage in p53-mutant cancer cell models

A clinical drug regimen for TAS102 and TAS102-talazoparib includes cycles of 5-days On-treatment plus a 9-day Off period ^37^. To explore the response dynamics, the cell-cycle analysis was performed on p53WT and p53-deficient cells treated with TAS102-talazoparib followed by drug washout. In p53WT cells, an increase in the G2 population was observed following the two-drug treatment and remained elevated up to 24h after drug washout, then followed by a gradual increase in the G1 population, with a small fraction of sub-G1 cells (**Fig. 4a**). In contrast, p53KO and p53R248W cells responded to TAS102-talazoparib with a prolonged G2-arrest even after drug withdrawal. Moreover, the sub-G1 population was increased at 48-72h after drug withdrawal, indicating cell death in p53-deficient cells (**Fig. 4b-c**).

**Figure 4.**
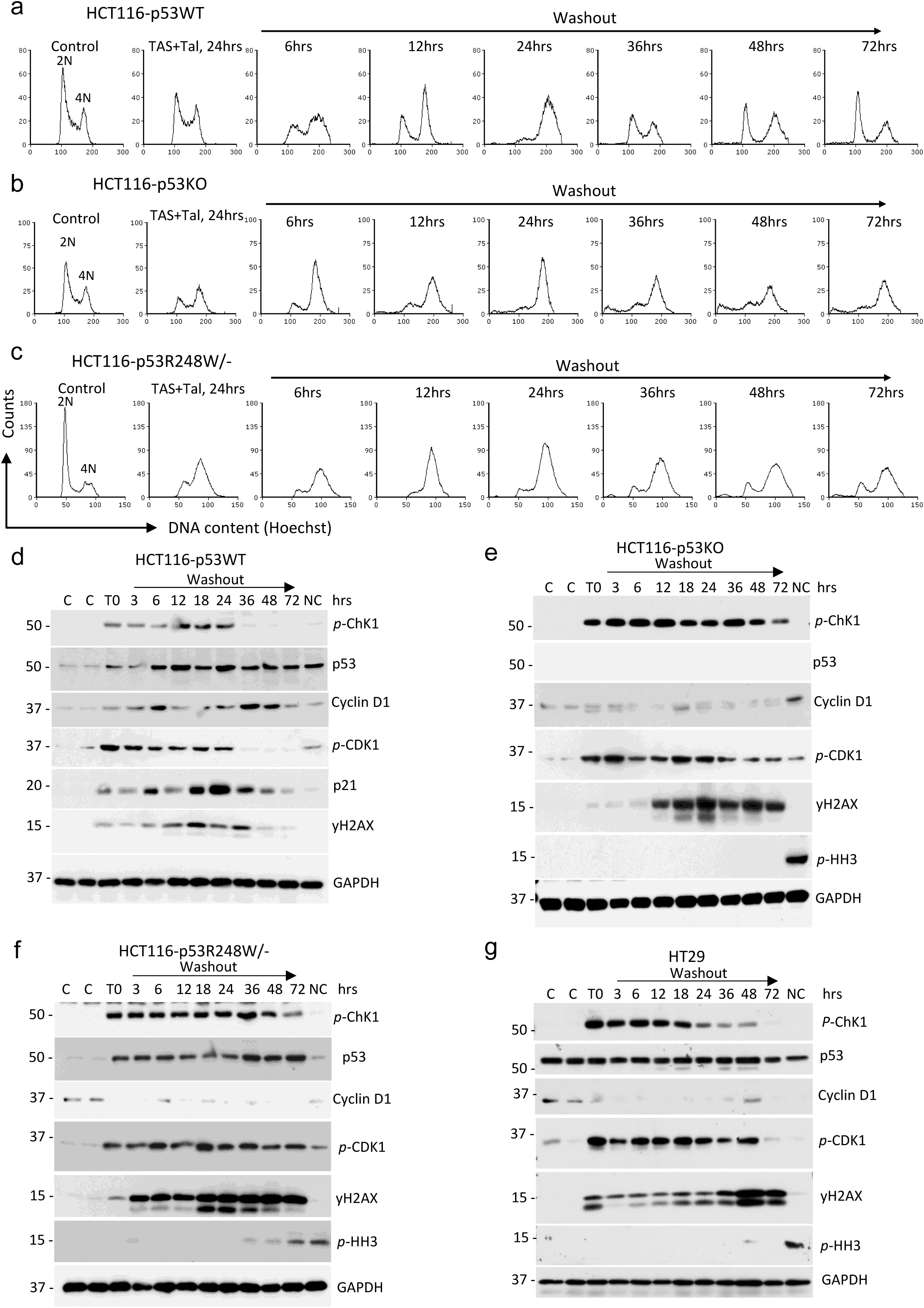
Combination treatment with TAS102 and talazoparib induces a prolonged G2 arrest in p53-mutant cancer cells. (**a-c**) Cell cycle analysis was performed on cell lines treated with vehicle control (C), 500nM TAS102 and 100nM talazoparib for 24 hours (**T0**) followed by incubation in drug-free medium. All experiments were done in at least two independent biological replicates. (**d-g**) Immunoblotting on whole-cell extracts from HCT116-p53WT, HCT116-p53KO, HCT116-p53^R248W/-^ and HT29 cell lines treated as described above (**a-c**). The experiments were repeated in at least two independent biological replicates and two technical replicates. Blots were probed with antibodies to phopsho-CHK1, p53, cyclin D1, phospho-CDK1, p21, phospho-H2AX (γH2AX), phospho-histone H3, and GAPDH as a loading control.

To verify the cell cycle effects and to assess DNA damage, the molecular markers of DDR and cell cycle were examined by immunoblotting in cells treated with TAS102-talazoparib for 24h followed by drug washout. In HCT116-p53WT cells, 24h-treatment with TAS102-talazoparib (T0) induced the markers of G2-checkpoint (phospho-CHK1 and phospho-CDK1) and DNA damage (γH2AX) (**Fig. 4d**). The levels of these markers decreased after 24hours of drug washout. Protein levels of p21/CDKN1A (a p53 target) followed this trend with a reduction at 36h (**Fig. 4d**). Cyclin D1 levels increased at 36h, indicating a recovery of cell cycle capacity (**Fig. 4d**). Together, these findings align with the cell cycle data (**Fig. 4a**). In contrast, HCT116-p53KO and HCT116-p53^R248W/-^ cells exhibited sustained phospho-CHK1, phospho-CDK1, and γH2AX levels after the TAS102-talazoparib treatment, indicating a prolonged G2-arrest with high DNA damage (**Fig. 4e-f**). Consistent with this, cyclin D1 protein levels remained low, not recovering within 72h (**Fig. 4e-f**). Comparable results were observed in HT29-p53^R273H^ cells (**Fig. 4g**), where a prolonged increase in DNA damage and G2 markers mirrored the data in HCT116 p53KO and p53R248W/-cell lines.

To validate the recovery of cell proliferation following treatment with TAS102-talazoparib for 24h and drug withdrawal, cell growth was examined using live-cell imaging. The analysis showed that p53WT non-tumor MCF10A cells quickly recovered growth capacity after drug washout, while growth recovery was delayed by ∼48h in p53WT HCT116 cells (**Suppl. Fig. 6a, b**). In contrast, p53KO and p53mut cancer cells did not regain the growth capacity within the 92h timeframe of the experiment (**Suppl. Fig. 6c-d**).

Together, these data showed that TAS102-talazoparib induced in p53-deficient cells a long G2-arrest and persistent DNA damage observed even after 72h of drug withdrawal, and a sizable proportion of p53mut cells underwent cell death within 48-72h (sub-G1 fraction). In contrast, p53WT cells were able to repair DNA and recover their cell cycle capacity within 48h of drug withdrawal.

### The dynamics of DNA repair following TAS102-talazoparib treatment in p53mut cells

To evaluate the dynamics of DNA damage repair, changes in the DSB markers γH2AX and RAD51 were examined in MCF10A (p53WT), DLD1 (p53^S241F^), and HT29 (p53^R273H^) cell lines using immunofluorescence. Cells were treated with TAS102-talazoparib for 24h followed by drug washout. Immediately after 24h-treatment (at T0), the levels and the foci formation of γH2AX and RAD51 were increased in all tested cell lines (**Fig. 5a, d, e, g, j, k, m, n**). In MCF10A cells, the nuclear area was increased by nearly 2-fold (from 130µm^2^ to 250µm^2^) (**Fig.5c, f**), indicating enlargement of the nucleus in a significant proportion of cells (about 77% with area of 200-580µm^2^, insert). Following drug withdrawal, the γH2AX and RAD51 signals and the nuclear sizes were decreased within 24-48h in MCF10A cells (**Figs. 5c-f**). These data show that DNA damage by TAS102-talazoparib was repaired, and cells completed the mitotic cell division. In contrast, the γH2AX/RAD51 signals were not reduced in p53mut DLD1 and HT29 cell lines within 72h of drug washout (**Fig. 5g-h, j-k, m-n**). The nuclear area was also increased at T0, and a sustained enlargement was observed at 24-48h of drug washout (**Fig. 5i, l, o**). These results indicate that p53mut cells replicate DNA but do not undergo cell division resulting in enlarged nucleus, and DSBs are not repaired up to 72h of drug washout. Together, these observations show that the TAS102-PARPi treatment can induce DNA breaks and G2-arrest in p53WT and p53mut cells. However, DNA repair and recovery of cell division were remarkably different in p53WT and p53mut cells. Non-tumor p53WT cells repaired DNA breaks and completed the mitotic cell division within 48h, whereas p53mut cells could not complete DNA repair and cell division, resulting in high levels of DNA breaks and G2-arrest even at 72hours of drug withdrawal. These observations are aligned with the cell cycle and cell growth data, which showed a lack of proliferation in p53mut cells within this timeframe (**Fig. 4, and Suppl. Fig.5**). Thus, TAS102-PARPi treatment induced high DSB levels and long G2-arrest in p53mut cancer cells, while p53WT cells repaired DNA and recovered cell cycle capacity.

**Figure 5.**
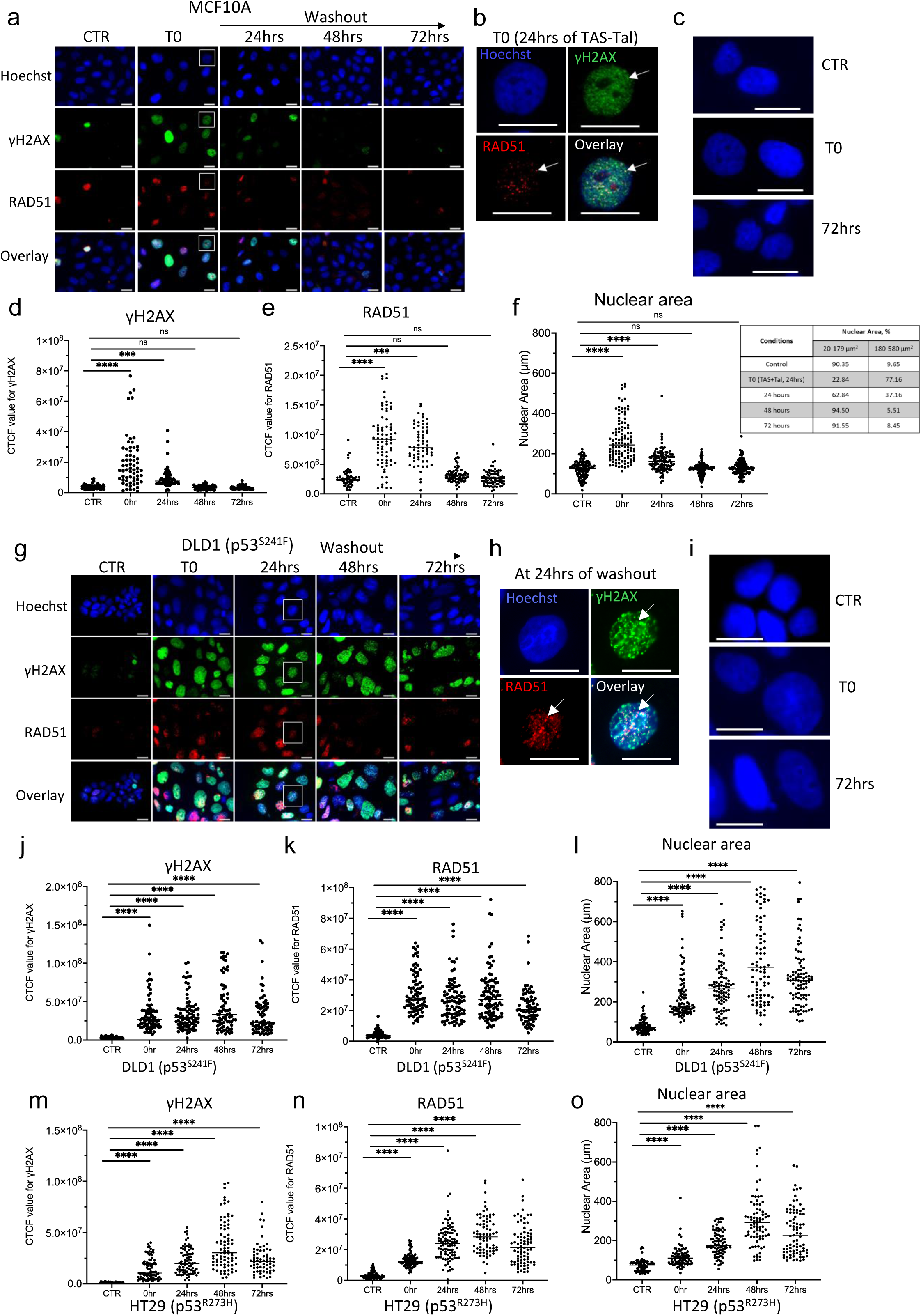
The dynamics of DNA damage repair in response to combined TAS102 and talazoparib treatment. (**a**) Immunofluorescence microscopy images of p53WT cells stained for γH2AX (green), a DNA damage marker, and RAD51 (red), a homologous recombination component, along with Hoechst 33342 (blue; nucleus). Cells were treated with vehicle-control (CTR) or 500nM TAS102 plus 100nM talazoparib (**T0**) for 24 hours, followed by drug withdrawal and the cells were allowed to recover. All images were taken using 100x lens; scale bar 10µm. (**b, h**) Enlarged images of regions highlighted in the main panels show the co-localization of γH2AX and RAD51 foci at T0 in MCF10A cells and at 24 hours in DLD1 cells. Arrows indicate the positions of these foci. (**c**) Enlarged images of Hoechst-stained cells at CTR, T0, and 72hours. (**d-e, j-k, m-n**) Quantitative analysis of corrected total cellular fluorescence (CTCF) for γH2AX (**d, j, m**) and RAD51 (**e, k, n**) was performed on a minimum 40 cells per treatment group. The graphs show representative median scattered dot plots. Comparisons were made using the two-way ANOVA (****, P<0.001). (**f, l, o**) The nuclear area was measured from Hoechst-stained cells, with at least 40 cells/group. All experiments were done at least two times.

### A sequential triple-drug approach using TAS102-PARPi treatment followed by blockade of WEE1 kinase elicits synergistic toxicity in p53mut cancer cells

TAS102-PARPi treatment induced DNA breaks and sustained G2-arrest in p53mut cells even after drug washout, whereas non-tumor p53WT cells recovered remarkably faster (**Figs. 4-5**). Next, we examined whether toxicity of TAS102-PARPi treatment in p53mut cells can be further enhanced by blockade of a G2-checkpoint kinase. Cells were treated with TAS102-talazoparib for 24h, followed by drug washout for 24h and subsequent incubation with WEE1i (MK1775) for 48h. Cytotoxicity assays showed that WEE1i did not synergize with TAS102-PARPi in p53WT MCF10A cells, ∼1.1-fold at 100nM WEE1i (**Fig. 6a, f**). In HCT116-p53WT cells, WEE1i only modestly enhanced the IC50 of TAS102-talazoparib (by ∼1.4-fold at 100nM WEE1i). In contrast, in HCT116-p53KO and HCT116-p53^R248W^ cell lines WEE1i at 100nM enhanced toxicity of TAS102-talazoparib by ∼3.6-fold and ∼2.5-fold, respectively (**Fig. 6b-d, f**). In HT29 (p53^R273H^) cells, the toxicity of TAS102-talazoparib was enhanced by ∼7-fold at 100nM WEE1i (**Fig. 6e-f**). Evaluation of drug interactions in this sequential regimen showed CI = 0.94 for MCF10A cells and CI = 0.57 for HCT116 cells (both p53WT), indicating no or modest drug interactions (**Fig. 6f**). In contrast, p53-deficient cells, HCT116-p53KO or p53mut cells (HCT116-p53^R248W^ and HT29-p53^R273H^), showed CI values ranging 0.26-0.42, indicating a synergistic interaction between TAS102-talazoparib and WEE1i (**Fig. 6f**).

**Figure 6.**
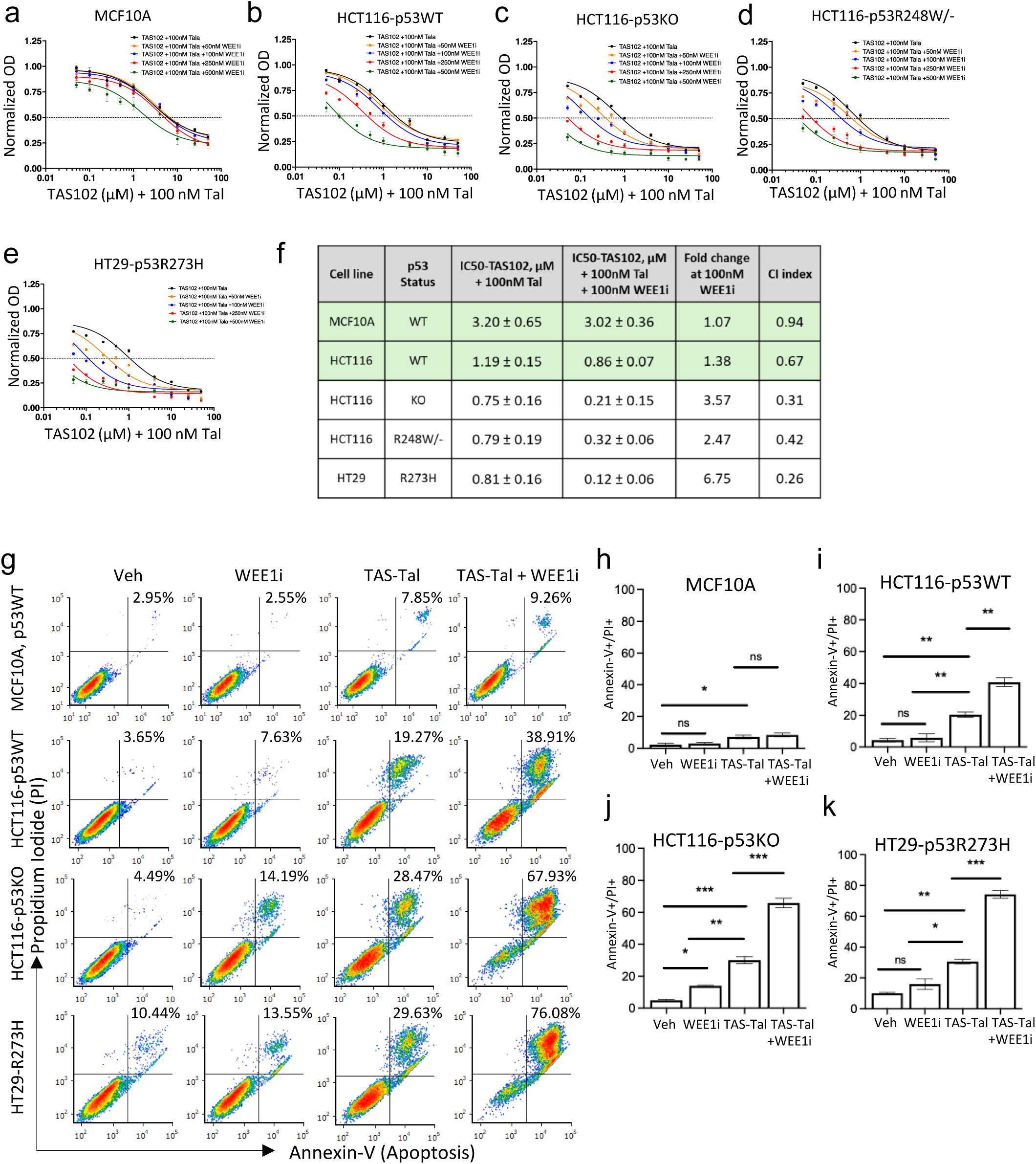
Sequential application of WEE1 inhibitor enhances the cytotoxicity of TAS102-talazoparib treatment in p53-deficient cancer cells. (**a-e**) Cytotoxicity curves for p53 wild-type (MCF10A and HCT116), and p53-deficient (HCT116-p53KO, HCT116-p53^R248W/-^, HT29) cells. Cells were treated with increasing concentrations of TAS102 plus 100nM talazoparib for 24 hours. After a 24-hour drug washout, cells were incubated with 50, 100, 250, or 500 nM of the WEE1 inhibitor (WEE1i) for an additional 72 hours. (**f**) Combination index (CI) analysis for the tested WEE1i concentrations. Data are presented as mean ± SEM from at least two independent biological replicates, each with six technical replicates. (**g**) Flow cytometry analysis of cells treated with vehicle control or a sequential regimen of 500 nM TAS102 plus 100 nM talazoparib, followed by 250 nM WEE1i. Cells were stained with propidum iodide and annexin-V. (**h-k**) Quantification of Annexin V and propidium iodide (PI) double-positive cells from two biological replicates. Statistical significance was determined using one-way ANOVA (*p < 0.05, **p < 0.01, ***p < 0.001, and ****p < 0.0001).

Examination of cell death, using flow cytometry with Annexin V and propidium iodide (PI) staining, showed that the sequential treatment of MCF10A cells with TAS102-PARPi followed by 100nM WEE1i resulted in ∼9% double-positive (annexin-V^+^/PI^+^) dead cells, compared to ∼8% dead cells by TAS102-PARPi treatment alone (**Fig. 6g-h**). In HCT116-p53WT cells, ∼19% of cells were double-positive after TAS102-talazoparib treatment and ∼39% after the sequential triple-drug combination (**Fig. 6g, i**). Approximately 29% of HCT116-p53KO cells were double-positive in response to TAS102-talazoparib, and cell death increased to ∼68% after triple-drug treatment (**Fig. 6g, j**). In HT29 cells, TAS102-talazoparib treatment resulted in ∼30% double-positive cells, whereas the sequential drug combination resulted in ∼76% double-positive dead cells (**Fig. 6g, k**). Similar effects of a sequential triple-drug treatment were observed in p53mut MDA-MB-231 cells compared to p53WT MCF10A cells (**Suppl. Fig. 7**). Together, these data showed a synergistic toxicity of TAS102-talazoparib treatment followed by WEE1i in p53mut cancer cells, whereas this sequential triple-drug regimen showed low toxicity in non-tumor p53WT cells.

### Anti-tumor efficacy of the sequential triple-drug combination

The therapeutic efficacy of the sequential combination of TAS102-talazoparib and WEE1i was examined in tumor xenograft models. The drug regimen included repeats of the two-week cycle with TAS102-talazoparib treatment in the first week followed by WEE1i treatment in the second week, based on the clinical schedule for TAS102 and TAS102-talazoparib ^37^. This two-week regimen was initially tested in xenografts of HT29 cells (p53^R273H^, BRAF^V600E^) in SCID mice. Treatment was initiated when tumors reached 100mm^3^ using 5-days-ON, 2-days-OFF schedule in four groups: vehicle, WEE1i (every week), TAS102-talazoparib (every week), and the triple-drug combination (a sequential two-week cycle). Treatment with WEE1i had a limited effect on tumor growth with tumor growth inhibition (TGI) of 18%, whereas TAS102-talazoparib was more effective, with a TGI of 43% (**Fig. 7a-b**), consistent with our previous studies^18^. Importantly, a sequential combination of TAS102-talazoparib and WEE1i showed greater tumor growth control than each treatment alone, with a TGI of 63% (**Fig. 7a-b**). The triple-drug regimen also improved survival to 39 days compared to 32 days for TAS102-talazoparib, and 24 days for WEE1i alone (**Fig. 7c**). Neither treatment affected animal weight compared to the control group (**Fig.7d**), indicating the absence of major toxic side effects.

**Figure 7.**
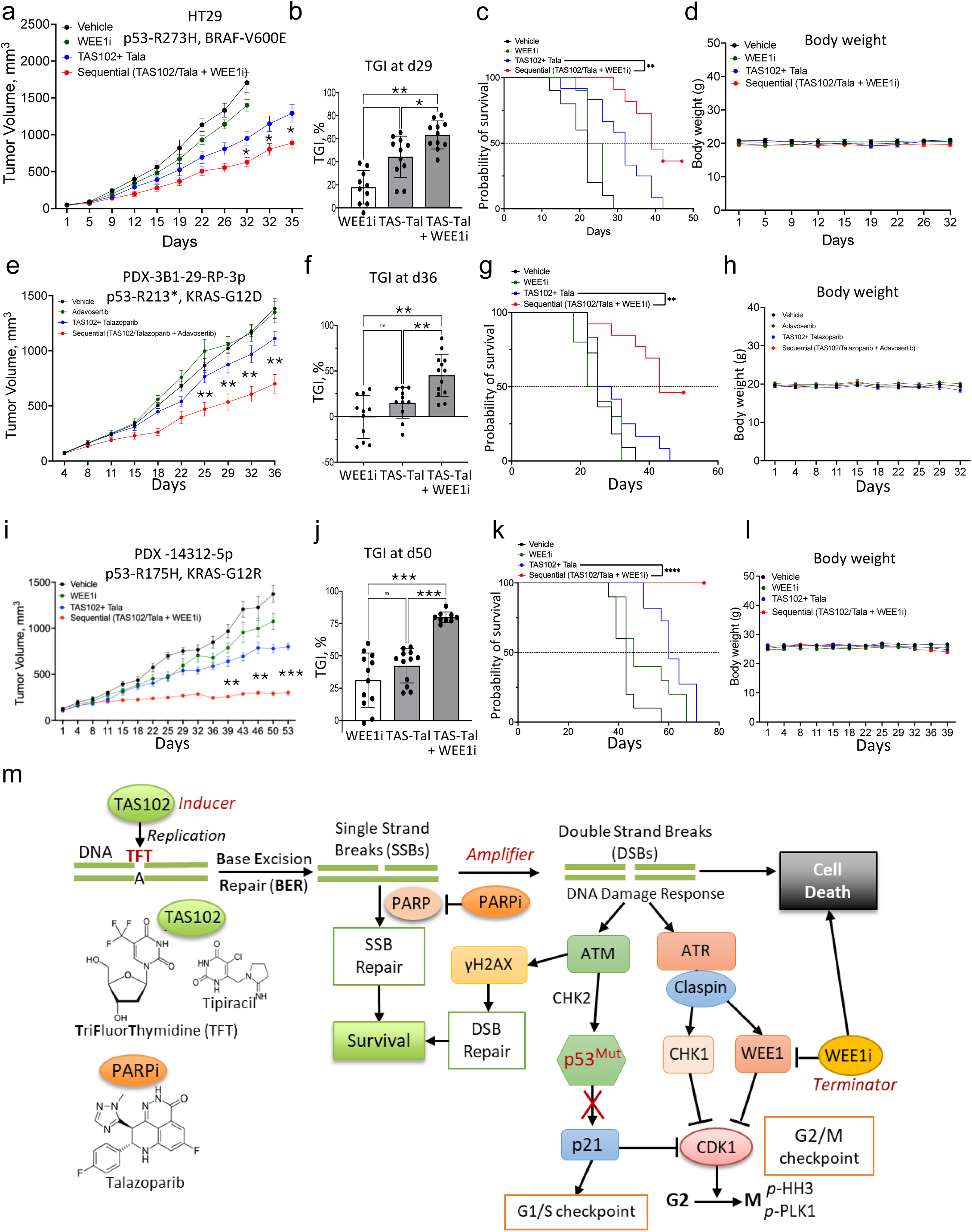
A sequential application of WEE1 inhibitor enhances the efficacy of TAS102-talazoparib treatment in p53 mutant cancer models. (**a-d**) SCID mice (10mice/group) were subcutaneously implanted with colon cancer HT29 cells. At tumor size of 100mm^3^, mice were randomized into four treatment groups: vehicle control, WEE1 inhibitor (WEE1i, MK1775, 60 mg/kg), a combination of TAS102 (50 mg/kg) and talazoparib (0.15 mg/kg), and a sequential regimen consisting of one week of TAS102/talazoparib followed by one week of the WEE1 inhibitor. All treatments were administered *via* oral gavage on a schedule of five days ON and two days OFF. Tumor growth was measured twice weekly, and statistical significance was determined using a two-way ANOVA (*p < 0.05). (**b**) Tumor growth inhibition (TGI) was calculated as TGI = 100% x [Vehicle − Treatment) / Vehicle]. Comparisons were made using a one-way ANOVA (**, P<0.01, ***, P<0.001, ****, P<0.001). (**c**) Survival was evaluated using Kaplan-Meier estimator based on time-to-arrive at 750 mm^3^ of tumor size. The comparison was made using the log-rank test. (**d**) Mouse body weights were measured twice per week over the course of study. (**e-h**) Patient-derived tumor xenografts (PDXs) from a patient with PDAC (PDAC-14312-4p) were subcutaneously implanted into SCID mice (10 mice/group). At 100 mm^3^, mice were randomized and treated as described above for HT29 xenografts. (**f**) Tumor growth inhibition in the PDX model was calculated as described in (**b**). Comparisons were made using a one-way ANOVA (**, P<0.01, ***, P<0.001, ****, P<0.001). (**g**) Survival was evaluated using Kaplan-Meier estimator based on time to arrive at 750 mm3 of tumor size. Group comparisons were performed with the log-rank test. (**h**) Mouse body weights were measured twice weekly over the course of study. (**i-l**) Patient-derived tumor xenografts (PDXs) from a colorectal cancer patient (PDX-3B1-29-RP-3p) were subcutaneously implanted into SCID mice (10 mice/group). Experiments and the analysis were done essentially as described for (**e-h**). (**m**) A model of the triple-drug drug strategy acting as an *inducer-amplifier-terminator* trio in p53 mutant cancers.

The efficacy of the triple-drug regimen was tested in a patient-derived xenograft (PDX) model (PDX-3B1-29-RP, p53^R213*^) established from a colorectal cancer biopsy within the clinical trial NCT04511039. The R213* truncating p53 mutation is commonly found in human tumors and exhibits pro-tumor effects ^38^. Treatment with TAS102-talazoparib reduced tumor growth by ∼20% and WEEi1 alone had no effect, whereas the sequential drug combination reduced tumor growth by ∼50% (**Fig. 7e-f**) and extended survival from 24 to 43 days (**Fig. 7g**). No major toxic side effects by the treatments were noted based on the examination of mouse weight (**Fig.7h**), the bone marrow histology, and peripheral cell blood counts (**Suppl. Fig. 8**). The efficacy was further examined in a pancreatic cancer PDX model (PDX-14312-5p) carrying p53^R175H^ and KRAS^G12R^ mutations. It was observed that WEE1i monotherapy modestly reduced tumor growth (∼30%), whereas TAS102-talazoparib was more effective in tumor control, with a TGI of ∼43% on day 50 (**Fig.7i-j**). The triple-drug regimen showed remarkably greater tumor control than either treatment alone (TGI ∼80%, **Fig.7i-j**) and extended survival to >75days, compared to ∼64 days by TAS102-talazoparib and 48 days by WEE1i (**Fig.7k**). No major toxicity signs were noted based on mouse weight (**Fig.7l**). Together, these data demonstrate that the triple-drug regimen combining the two-drug TAS102-PARPi treatment followed by WEE1i is effective against p53mut cancer models and the regimen is well tolerated in mice.

## DISCUSSION

Treatment of p53mut cancers remains a significant clinical challenge mainly due to lack of effective therapeutic options ^9, 10^. Here, we introduce a triple-drug strategy for p53mut cancers combining two-drug TAS102-PARPi treatment with blockade of the G2-checkpoint. TAS102-PARPi treatment acts as an *inducer-amplifier* pair to induce DNA double-strand breaks (DSBs) and G2-arrest selectively in p53mut cancer cells. Subsequent blockade of the G2-checkpoint kinase WEE1 releases G2-arrested cells into mitosis leading to cell death, acting as a *terminator*. We found that delaying WEE1i application until after TAS102-PARPi treatment allows normal p53WT cells to repair their DNA, thus reducing the impact on normal cells (**Figs. 5-6**). This sequential triple-drug approach is a promising therapeutic strategy for treatment of p53mut cancers as it showed high efficacy in preclinical cancer models and was well tolerated in mice.

Prior work has shown that TAS102-PARPi treatment induces high levels of DNA damage in p53mut cancer cells, and this effect was linked to a compromised G1/S checkpoint ^17, 18^. Thymidine (dT) analogs, such as trifluorothymidine (TFT, a component of TAS102) or ethynyl-deoxyuridine (EdU), are incorporated into DNA during replication^17, 18^. These dT-analogs also inhibit thymidylate synthase^39^, leading to the incorporation of uridine residues into DNA^17, 18, 40^. The incorporated analogs are removed from DNA by base-excision repair (BER) resulting in single-strand breaks (SSBs) ^17, 18, 40^, which are repaired *via* a mechanism involving PARP1^41^. PARP1 inhibitors (such as talazoparib) compromise SSB repair, increasing levels of more lethal DSBs that further activate DDR signaling mediated by ATM and ATR kinases. In p53WT cells, DDR signaling mediated by ATM activates the p53-p21 axis resulting in G1-checkpoint activation ^18^. p21/CDKN1A inhibits CDK2 and CDK4/6 leading to cell cycle arrest in G1/S and/or inhibit CDK1-cyclin B1 contributing to G2-arrest ^42–44^. This effect of TAS102-PARPi treatment is consistent with the role of ATM-CHK2-p53 signaling in DNA damage response ^42,43^. In p53mut cells with compromised G1/S checkpoint, TAS102-PARPi treatment resulted in high DSBs and activation of the G2-checkpoint mediated by ATR (this study and ^18^).

The distinct effects of TAS102-PARPi treatment on p53WT and p53-deficient cells were also reflected in the transcriptomic program. In p53WT cells, TAS102-PARPi reduced expression of genes involved in the G1/S transition (e.g., E2F1, E2F3), indicative of G1-arrest. However, in p53-deficient cells, TAS102-PARPi increased expression of the G1/S transition factors (E2F1, E2F3) and S/G2-phase genes (e.g., CCNE1, CCNE2, CDT1), consistent with the lack of G1/S checkpoint in these cells. Moreover, TAS102-PARPi treatment reduced expression of M-phase genes (i.e., AURKA, BUB1, CDC20, PLK1), consistent with G2-arrest in p53-deficient cells. Thus, TAS102-PARPi treatment had distinct effects on p53WT and p53mut cells in the global gene expression program related to cell cycle control.

Transcriptome analysis also revealed striking upregulation of HR-related DSB repair genes (BRCA1, BRCA2, RAD51, RECQL, RBBP6) in p53-deficient cells. This distinct effect of TAS102-PARPi treatment is likely to reflect DNA repair at the G2-checkpoint controlled by ATR kinase ^45^. ATM may also contribute to DNA repair by acting through H2AX (γH2AX), RAD51, and BRCA1/2, thus promoting homologous recombination (HR) ^46–48^. Thus, in addition to G2-arrest, TAS102-PARPi treatment upregulated HR-mediated DNA repair.

Cytotoxicity studies showed a substantial dichotomy in the consequences of ATM and ATR inhibition. Blockade of ATM reduced sensitivity to TAS102-talazoparib treatment in all tested lines cells, independent of the p53 status. This effect of ATM blockade is likely related to the function of ATM in the G1/S checkpoint ^23^ or replication fork recovery after DNA repair ^21, 22^. In contrast, ATR blockade greatly enhanced cytotoxicity of TAS102-talazoparib in p53mut cells, and this effect is related to transition to mitosis with high DNA damage (elevated phospho-H3 and γH2AX levels). These results are consistent with the role of ATR in activation of the G2/M checkpoint and in the control of replication completion before mitosis ^24^. ATR can also control an intrinsic S/G2 checkpoint, and the ATR blockade may deregulate the S/G2 transition allowing early mitosis with under-replicated DNA and DNA damage ^45^. Thus, ATR plays a critical role in activation and control of the G2-checkpoint in p53mut cells treated with TAS102-PARPi, making ATR a potential target to further enhance efficacy of TAS102-PARPi treatment.

To induce the G2 checkpoint, ATR activates downstream effector kinases, such as CHK1 and WEE1, which inactivate the CDC25-CDK1 axis that controls transition to mitosis ^21, 25–30^. Here, we found that CHK1 blockade in combination with TAS102-PARPi treatment was cytotoxic in both non-tumor p53WT and p53mut cancer cells (**Fig. 3**). This effect may reflect CHK1 role in S-phase and explain high toxicities of CHK1 inhibitors observed in clinical trials, leading to discontinuation of early trials ^31, 32^. In contrast, WEE1 blockade enhanced toxicity of TAS102-talazoparib preferentially in p53mut cells, while showing limited toxicity in non-tumor models (**Fig.3**). This effect is consistent with the tolerable toxicities of WEE1i observed in early-phase clinical trials ^33, 34^.

The drug response studies showed that the selectivity and efficacy of the drug combinations can be enhanced in a sequential triple-drug regimen. TAS102-talazoparib treatment of p53mut cancer cells resulted in prolonged G2-arrest and persistent DNA damage even at 72h after drug washout with a sizable amount of cell death (**Figs.4-5**). In contrast, p53WT cells were able to repair DNA and re-enter the cell cycle within 48h of drug withdrawal. Moreover, a 24h-delay in WEE1i treatment after TAS102-talazoparib washout was still highly effective against p53mut cells but was markedly less toxic to normal cells (**Fig.6**). This sequential triple-drug regimen (TAS102-talazoparib plus 24h-delayed WEE1i) resulted in massive 80% cell death in p53mut cancer cells, whereas cell death was not increased in non-tumor cells.

A sequential triple-drug regimen combining TAS102-talazoparib and WEE1i showed robust anti-tumor effects in preclinical animal models (**Fig. 7**). Tumor growth was reduced by 63% in the CRC HT29 tumor model and ∼80% in the PDAC PDX-14312 model (p53^R175H^) (**Fig.7**). Further, the triple-drug treatment was also highly effective in reducing tumor growth in CRC PDX model (∼55%). No major toxic side effects were observed during assessment of mouse weight, bone marrow, and peripheral blood.

In summary, this study presents a sequential triple-drug strategy combining TAS102-PARPi treatment with delayed application of a G2-kinase inhibitor (WEE1i), which together work as an *inducer-amplifier-terminator trio* (**Fig.7m**). The TAS102-talazoparib regimen was well tolerated in patients with refractory CRC and showed promising median progression-free survival (mPFS) of 5.7 months (NCT04511039) ^37^. This study showed that a sequential G2-kinase blockade with WEE1i strongly enhances efficacy of TAS102-talazoparib in p53mut cancers. This sequential triple-drug strategy is a promising approach for management of patients with p53mut cancers.

### Limitations of the Study

This study demonstrated the increased efficacy of TAS102-PARPi treatment in combination with a subsequent administration of WEE1i in p53mut CRC and PDAC models. Although, prior work has shown that TAS102-PARPi treatment can block breast cancer metastases ^17^, the current studydid not directly assess the effect of the combinatorial drug treatment on metastatic burden (e.g., liver) in PDAC and CRC *in vivo* models. This question is the subject of future research.

## AUTHOR CONTRIBUTIONS

A.V.B. conceived the idea; A.V.B., R.M.S., J.Z., and M.M.A. designed the experiments; J.Z., M.M.A., A.V.B, M.N.N and B.G. performed the experiments and analyzed the data; A.V.B. and C.F. analyzed and interpreted the data; T.M. and R.M.S. helped in the interpretation of the data; B.G., B.F., S.C. helped with PDX generation; A.V.B., C.F., T.M., J.Z., M.M.A., R.M.S. and R.I. contributed to preparing and writing the paper.

## ABBREVIATIONS

BER: base excision repair
CRC: colorectal adenocarcinoma
DSB: double-strand break
EdUrd: 5-ethynyl-2’-deoxyrudine
TFT: trifluorothymidine
HR: homological recombination
MMR: mismatch repair
mPFS: median progression-free survival
OS: overall survival
PDAC: pancreatic ductal adenocarcinoma
SSB: single-strand break

## ACKNOWLEDGEMENT

We gratefully acknowledge the generous help from Flow and Image Cytometry Facility, the Pathology Resource Network, Preclinical Imaging Services, Experimental Tumor Model Shared Resource, and Laboratory Animal Resource (LAR), Ninfa L. Straubinger for technical assistance with the PDAC models. We thank Drs. Bert Vogelstein and Alessandro Carugo for their generous gifts of cell lines, Drs. Dean Tang and David Goodrich for discussion of the manuscript. This work supported by NIH R21CA259719 and DoD BC220542 (to AVB), NIH R37CA282430 (to CF and AVB), Roswell Park Alliance Foundation (to AVB and CF), NIH R01CA198096 to R.M.S., NIH grant AI164081 (to TM), in part by NIH R25CA181003, the Deanship of Scientific Research at Northern Border University, Arar, KSA the project number “NBU-SAFIR-2024 to (MMA), and the Roswell Park Comprehensive Cancer Center Support Grant, P30CA016056. Research reported in this publication was supported by the National Institutes of Health Office of Research Infrastructure Programs under award number S10OD024973 ^49^. The content is solely the responsibility of the authors and does not necessarily represent the official views of the National Institutes of Health. Computational support was provided by the Center for Computational Research at the University at Buffalo ^50^.

## ETHICS STATEMENT

The study protocol was approved by the Institute Animal Care and Use Committee (IACUC). The facility was certified by the American Association for Accreditation of Laboratory Animal Care (AAALAC) and in accordance with the current regulations and standards of the US Department of Agriculture and the US Department of Health and Human Services.

## MATERIALS AVAILABILITY

Patient-derived xenograft colorectal cancer models are available from the lead contact (A.B.) under a Material Transfer Agreement.

## LEAD CONTACT

Further information and requests for resources and reagents should be directed to and will be fulfilled by the lead contact, prof. Andrei Bakin

## KEY RESOURCES TABLE

| REAGENT or RESOURCE | SOURCE | IDENTIFIER |
| --- | --- | --- |
| **Antibodies** |  |  |
| anti-p53 (Mouse Monoclonal antibody, Clone do-1) | Santa Cruz Biotechnology | Cat# sc-126;<br>RRID:AB_628082 |
| anti-p21 (CDKN1A), F-5 mouse monoclonal | Santa Cruz Biotechnology | Cat# sc-6246;<br>RRID:AB_628073 |
| anti-phospho-Ser15 p53 (rabbit, polyclonal) | Cell Signaling Technology | Cat# 9284;<br>RRID:AB_331464 |
| anti-PARP1 (mouse monoclonal antibody, Clone 5a5) | Santa Cruz Biotechnology | Cat# sc-56197;<br>RRID:AB_630080 |
| anti-phospho-Ser139 H2AX (γH2AX; rabbit polyclonal) | Abcam | Cat# ab-11174;<br>RRID:AB_297813 |
| anti-Thymidylate Synthase (TS106) | Millipore | Cat# MAB4130,<br>RRID:AB_2210729 |
| anti-phospho-Tyr15-Cdc2 | Santa Cruz | Cat# sc-136014,<br>RRID:AB_2016914 |
| anti-phospho-Chk1 (Ser325) | Cell signaling | Cat# 2348,<br>RRID:AB_331212 |
| anti-phospho HH3 (Ser10) | Cell signaling | Cat# 31261,<br>RRID:AB_3076725 |
| anti-Cyclin A2 | Invitrogen | Cat# MA1-180,<br>RRID:AB_2633305 |
| anti-bromodeoxyuridine, anti-BrdU | Dako | Cat# M0744,<br>RRID:AB_10013660 |
| anti-cleaved Caspase-3 (Asp175) | Cell signaling | Cat# 9661,<br>RRID:AB_2341188 |
| Rabbit anti-gamma-H2AX (phospho-Ser139) Antibody | Bethyl | Cat# A300-081A,<br>RRID:AB_203288 |
| anti-Human Ki67/MKI67 Antibody | R&D Systems | Cat#: IC7617S,<br>RRID:AB_2889352 |
| anti-Rad51 antibody | Abcam | Cat# ab133534,<br>RRID:AB_2722613 |
| anti-phospho-Ser139 H2AX (γH2AX; mouse mAb, JBW301) | Millipore | Cat# 05-636;<br>RRID:AB_309864 |
| anti-BrdU antibody (3D4), FITC conjugate | Biolegend | Cat#364104;<br>RRID:AB_2564481 |
| Goat Anti-Rabbit IgG (H L)-HRP Conjugate antibody | Bio-Rad | Cat# 170-6515;<br>RRID:AB_11125142 |
| Goat Anti-Mouse IgG (H L)-HRP Conjugate antibody | Bio-Rad | Cat# 170-6516;<br>RRID:AB_11125547 |
| **Biological samples: Patient Derived Xenografts (PDXs)** |  |  |
| PDAC PDX-14312-4p | Dr. Robert Straubinger (University at Buffalo) | SUNY at Buffalo |
| CRC PDX-3B1-29-RP-3p | Dr. Andrei Bakin (Roswell Park) | PDX-3B1-29-RP |
| Chemicals, peptides, and recombinant proteins |  |  |
| Talazoparib (Synonyms: BMN-673) | Adooq | Cat# A15533 |
| TAS102 (trifluorothymidine (trifluridine)/tipiracil HCl mix) | Adooq | Cat# A11243 |
| 5-ethynyl-2'-deoxyuridine (EdUrd) | Lumiprobe | Cat# 20540 |
| Cy3-azide | Lumiprobe | Cat#B1030 |
| CellTrace Violet Dye | Invitrogen | Cat#C34557 |
| AFDye 488 Azide | Click Chemistry Tools, Scottsdale, AZ | Cat#1275-1 |
| THPTA (tris-hydroxypropyltriazolymethylamine) | Click Chemistry Tools, Scottsdale, AZ | Cat#1010-500 |
| RNase A (10 mg/mL) | Thermo Fisher Scientific | Cat#EN0531 |
| D-luciferin | Gold Biotechnology, St Louis | LUCK-1G |
| Copper sulfate, Cu(II)SO<sub>4</sub> | Sigma-Aldrich | Cat# 61230 |
| Hydroxyurea | Sigma-Aldrich | Cat#55291 |
| (2-Hydroxypropyl)-β-cyclodextrin (HPCD; Cavasol) | Sigma-Aldrich | Cat#778907 |
| Bovine serum albumin (BSA) | Sigma-Aldrich | Cat#A2153 |
| Dimethyl sulfoxide (DMSO) | Sigma-Aldrich | Cat#D8418 |
| Boric acid | Sigma-Aldrich | Cat#B6768 |
| Propidium iodide | Sigma-Aldrich | Cat#P4170 |
| Hoechst 33342 | Sigma-Aldrich | Cat#B2261 |
| Methylene Blue hydrate | Sigma-Aldrich | Cat#M9140-100G |
| Paraformaldehyde | Sigma-Aldrich | Cat#158127 |
| Dialyzed FBS (fetal bovine serum) | Gibco | Cat#A33820 |
| ECL Western Blotting Substrate, Pierce | Thermo Fisher Scientific | Cat# 32106 |
| SuperSignal™ West Atto Ultimate Sensitivity Substrate | Thermo Fisher Scientific | Cat#A38555 |
| SDS (Sodium Dodecyl Sulfate) | Bio-Rad | Cat#1610301 |
| DC™ Protein Assay Kit I | Bio-Rad | Cat#5000111 |
| Tween 20, 100% Nonionic Detergent | Bio-Rad | Cat#1706531 |
| Matrigel Matrix GFR | VWR International | Cat# 47743-720 |
| Critical commercial assays |  |  |
| Caspase-Glo® 3/7 Assay System | Promega | Cat#G8090 |
| Deposited data |  |  |
| Gene Expression and Mutation Counts | Pan-CANCER-TCGA | Cbioportal.org |
| Experimental models: Cell lines |  |  |
| HCT-116 (p53-WT) | Dr. Bert Vogelstein (Johns Hopkins) | RRID:CVCL_0291 |
| HCT-116-p53-KO; HCT-116 TP53(-/-); HCT116-p53-null | Dr. Bert Vogelstein (Johns Hopkins) | RRID:CVCL_HD97 |
| HCT-116-p53-R248W/- | Dr. Bert Vogelstein<br>(Johns Hopkins) | RRID:CVCL_RJ68 |
| RKO | Dr. Alessandro Carugo<br>(IRBM European<br>School of Molecular<br>Medicine, Lazio, Italia) | RRID:CVCL_0504 |
| RKO-p53(-/-); RKO-p53KO; RKO-p53-null | Dr. Alessandro Carugo<br>(IRBM ESMM, Lazio,<br>Italia) |  |
| HT29; HT-29 | ATCC | RRID:CVCL_0320 |
| MIAPACA2; MiaPaCa-2 | ATCC | RRID:CVCL_0428 |
| MCF10A | ATCC | RRID:CVCL_0598 |
| MDA-MB-231 | ATCC | RRID:CVCL_0062 |
| HPNE; hTERT-Human Pancreatic Nestin-Expressing cells | ATCC | RRID:CVCL_C466 |
| DLD1 | ATCC | RRID:CVCL_0248 |
| PANC-1 | ATCC | RRID:CVCL_0480 |
| Experimental models: Organisms/strains |  |  |
| Mouse: SCID/CB17, Balb/c background | Roswell Park Colony |  |
| Mouse: NOD.Cg-Prkdcscid Il2rgtm1Wjl/SzJ (NSG) mice,<br>Balb/c background | Jackson Laboratory |  |
| Software and algorithms |  |  |
| GraphPad PRISM 10.1.2 | GraphPad Software | https://www.graphpad.com/ |
| R/RStudio v4.0.4 | The R Foundation | https://www.r-project.org/ |
| FCS Express 7 (Version 7.04.0016) | De Novo software | FCS Express Flow<br>Cytometry Software<br>- De Novo Software |
| Image J | NIH | https://imagej.net/software/imagej/ |
| HeskaView Integrated Software (version 2.5.2) | Heska | Element HT5 - Heska |
| Aperio ImageScope (version 12.4.0.5043) | Leica Biosystems |  |
| MetaVue imaging software (Version 7.7.3) | Molecular Devices |  |
| FACSDiva (Version 6.1.3) | Beckman Coulter |  |
| Other |  |  |
| Nikon TE2000-E inverted microscope | Nikon, USA |  |
| CoolSNAP HQ (CCD camera) | Photometrics, USA |  |
| LSRFortessa Cytometer | BD Biosciences, USA |  |
| VERITAS microplate luminometer | Promega, USA |  |
| Nanodrop | Thermo Fisher, USA |  |

## Notes

### Competing Interest Statement

The authors have declared no competing interest.

### Summary of Updates

The revision includes changes in the Summary/Abstract. These changes are made to better outline novelty and impact of the study.

