## Supplemental Figures for "A Sequential Triple-Drug Strategy for Selective Targeting of p53-Mutant Cancers"

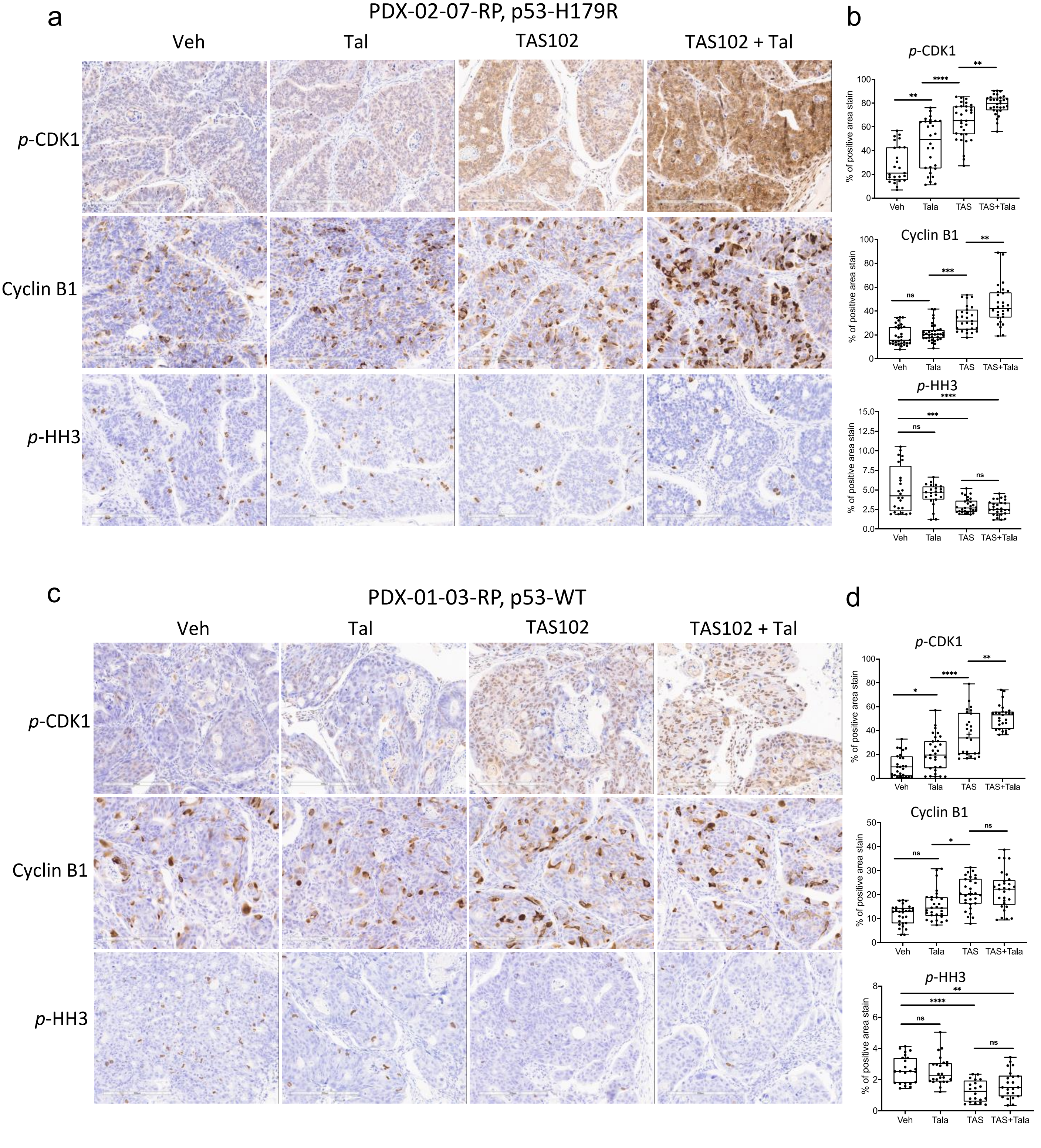


**Suppl. Fig.1. Immunohistochemistry (IHC) analysis of G2/M cell cycle markers in colorectal cancer PDX tissues treated with TAS102-talazoparib.** (**a, c**) Colorectal cancer PDX tissues were stained for phospho-CDK1, cyclin B1 and phospho-histone H3. (**b, d**) Quantifications were done using three tumors per group, with at least 25 fields per stain. Statistical comparisons were performed using one-way ANOVA (*p < 0.05, **p < 0.01, ***p < 0.001, and ****p < 0.0001). The analysis was done using three samples per condition.

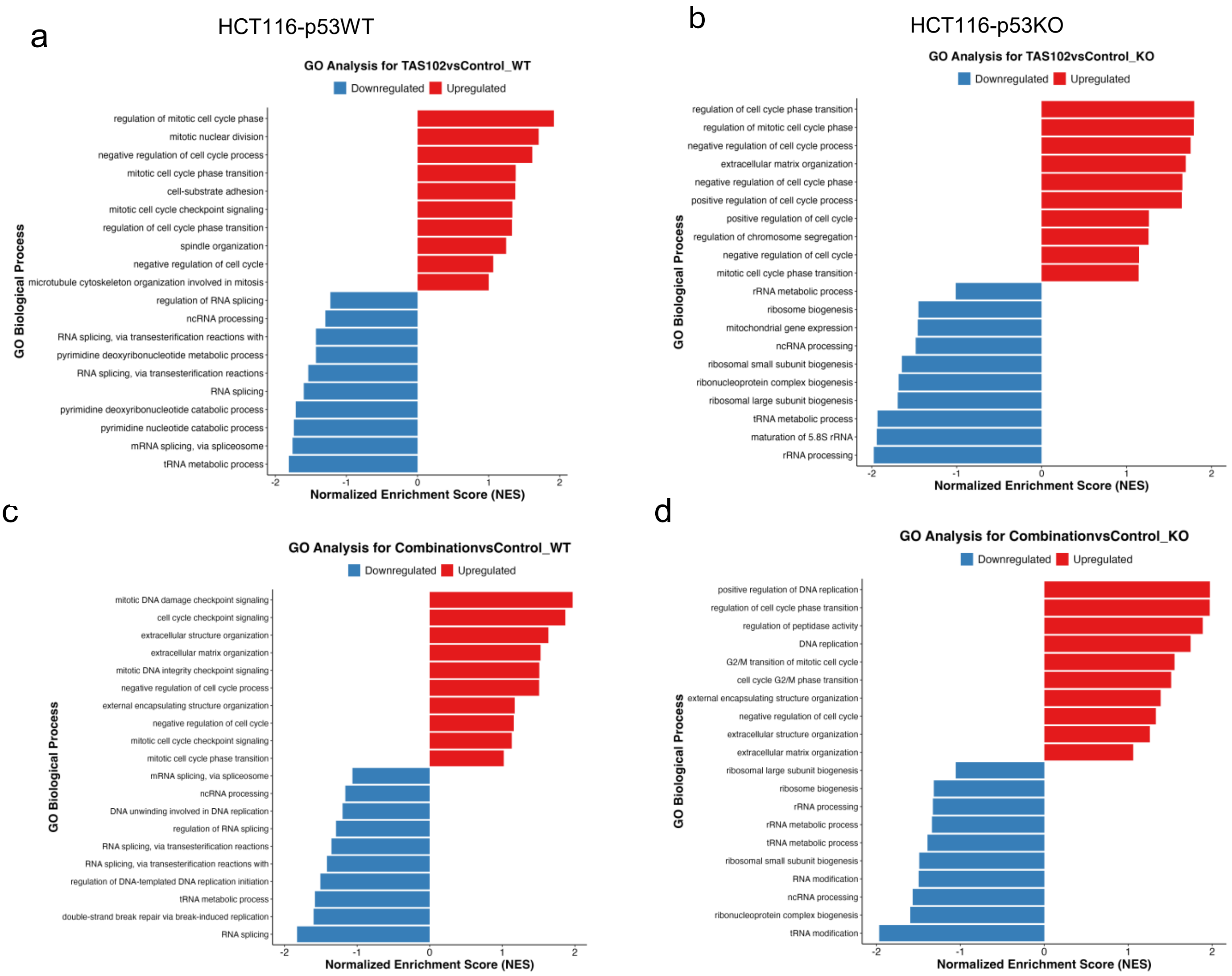


**Suppl. Fig.2. Pathway enrichment analysis**. (**a-d**) Gene Ontology (GO) analysis of the top 10 biological processes upregulated (red bars) and downregulated (blue bars) in HCT116 p53WT and HCT116 p53KO cells treated with TAS102 *vs*. Control-Vehicle (**a-b**) and the TAS102-talazoparib combination *vs*. Control-Vehicle (**c-d**). Pathways were selected based on adjusted p-values, and the NES was calculated by ranking all genes based on their differential expression, calculating the Enrichment Score (ES) for each gene set, and normalizing the ES by dividing it by the mean ES values from random permutations.

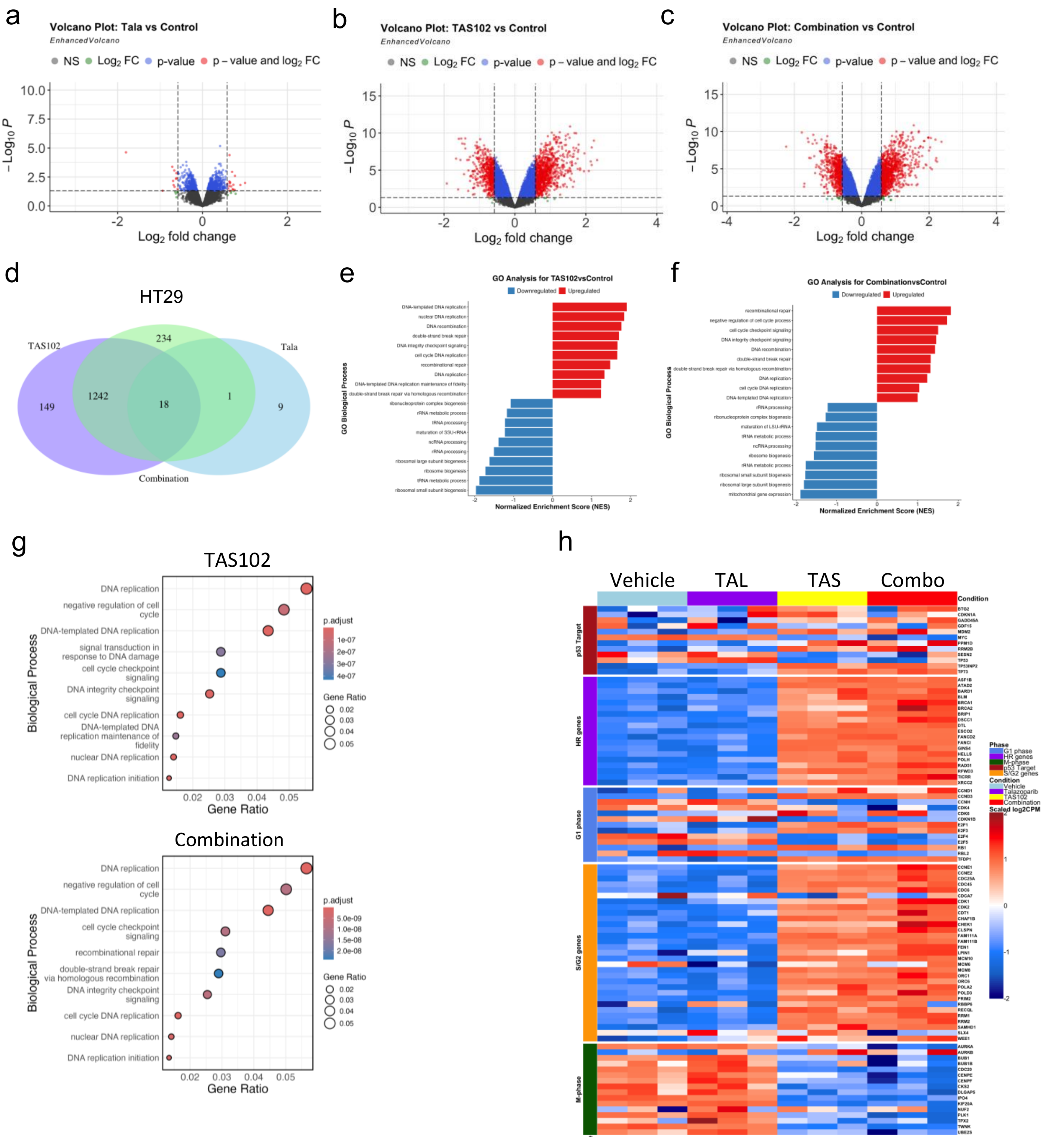

**Suppl. Fig.3. Differential Gene Expression in HT29 cells Upon Treatment.** (**a-c**) Volcano plots of differential gene expression (DGE) in HT29 cells treated for 24 hours with vehicle-control, 100 nM talazoparib (Tal), 500 nM TAS102, and TAS102 plus talazoparib in combination. (**d**) Venn diagram showing the overlap of differentially expressed genes in HT29 cells. (**e-g**) Gene Ontology (GO) analysis of the top 10 biological processes upregulated (red bars) and downregulated (blue bars) in HT29 cells treated with TAS102 *vs*. Control-vehicle (**e**) and the TAS102-Tala combination vs. Control-vehicle (**f**). Pathways are shown based on adjusted p-values. The normalized enrichment score (NES) was calculated by ranking all genes based on their differential expression, calculating the Enrichment Score (ES) for each gene set, and normalizing the ES by dividing it by the mean ES values from random permutations. (**h**) Heatmaps of differential expression in HT29 treated with vehicle-control, 100 nM talazoparib (Tal), 500 nM TAS102 (TAS), or their combination for 24 hours. Cell-cycle gene lists were derived from the Cyclebase 3.0 database. RNAseq analysis was done on triplicate samples.

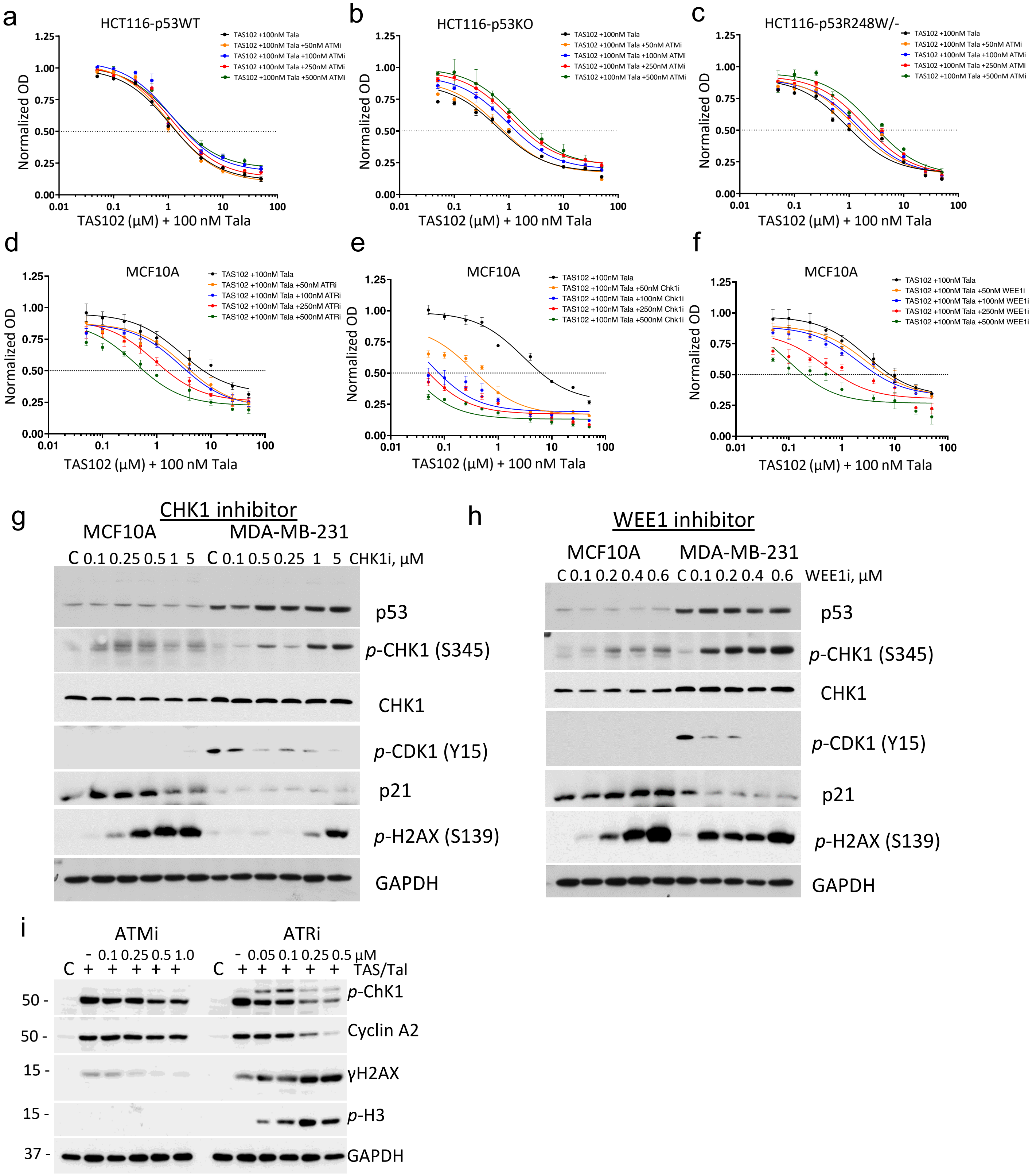


**Suppl. Fig. 4. Effects of G2 kinase inhibitors.** (**a-c**) Cytotoxicity of ATM inhibitor (ATMi) in combination with TAS102-talazoparib regimen. Cytotoxicity curves in CRC cell lines treated for 72hrs with TAS102 (μM) plus 100 nM talazoparib in combination with ATMi, KU55933, at 50nM, 100nM, 250nM, and 500nM. (**d-f**) Effects of G2 kinase inhibitors in non-tumor p53WT cells. Cytotoxicity curves in non-tumor MCF10A p53WT cells treated for 72hrs with TAS102 (μM) plus 100 nM talazoparib in combination with kinase inhibitors: (**d**) ATR inhibitor (ATRi), AZD6738, at 50nM, 100nM, 250nM, and 500nM; (**e**) CHK1i, AZD7762, at 50nM, 100nM, 250nM, and 500nM; and (**f**) WEE1i, MK1775, at 50nM, 100nM, 250nM, and 500nM. (**g-i**) Immunoblot analysis of effects of G2 kinase inhibitors. Cells were treated with various concentrations of WEE1 inhibitor (MK1775) or CHK1 inhibitor (AZD7762) for 24 hours and probed with antibodies to indicated targets. (**i**) HT29 CRC cells were treated with TAS102-talazoparib plus various concentrations of ATMi or ATRi. All experiments were repeated at least 2 times.

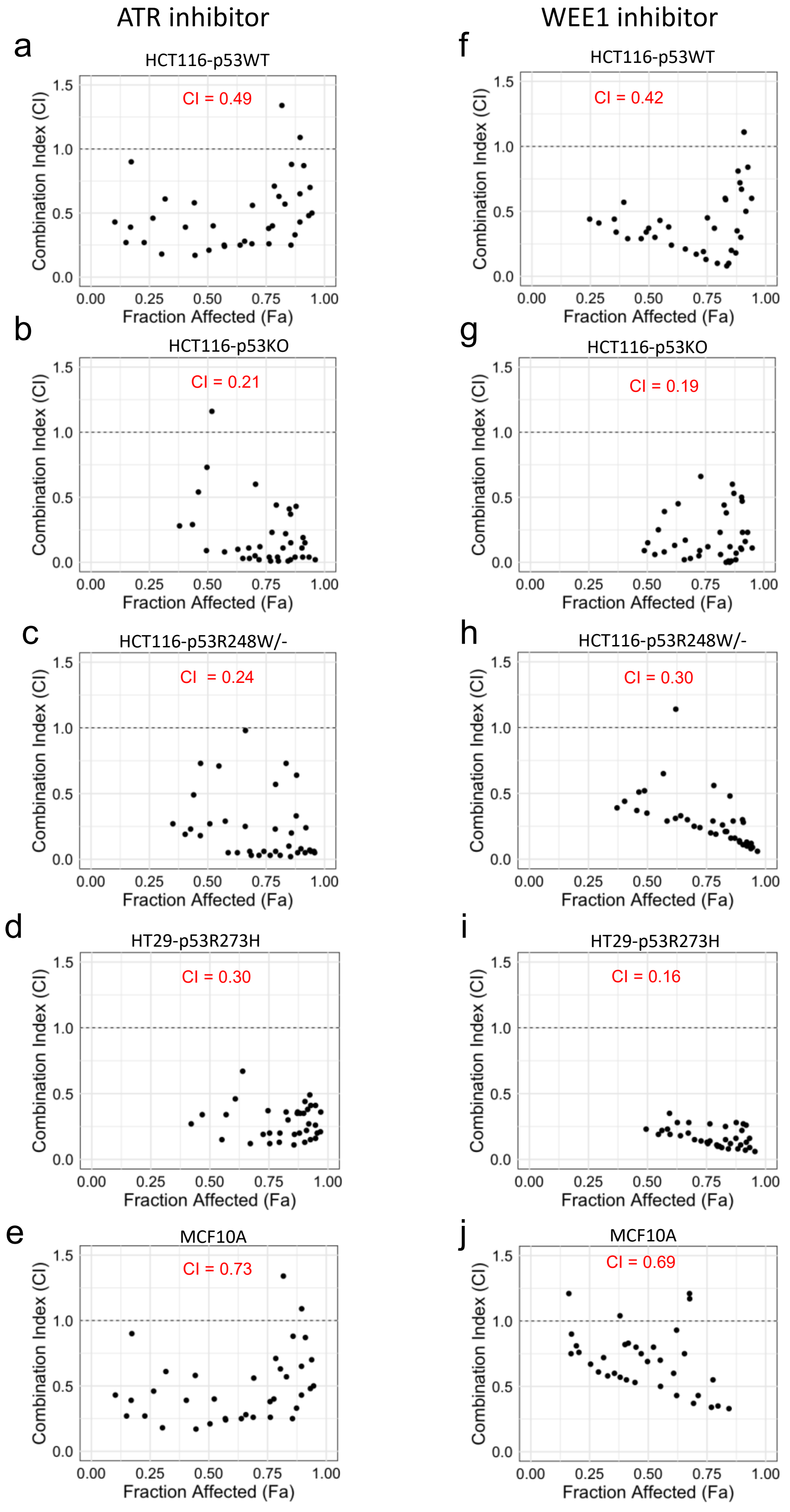


**Suppl. Fig. 5. Evaluation of drug interactions.** The Combination Index (**CI**) was calculated using the Loewe Additivity Model of TAS102 + talazoparib combination with ATRi (**a-e**) and WEE1i (**f-j**). Each dot in the scatter plot represents the interaction between the drug combination at specific concentrations, plotted against the fraction affected (**Fa**). The CI values are distributed below, at, or above the threshold of 1, indicating synergy (CI < 1), additivity (CI = 1), or antagonism (CI > 1). All experiments were repeated at least 2 times.

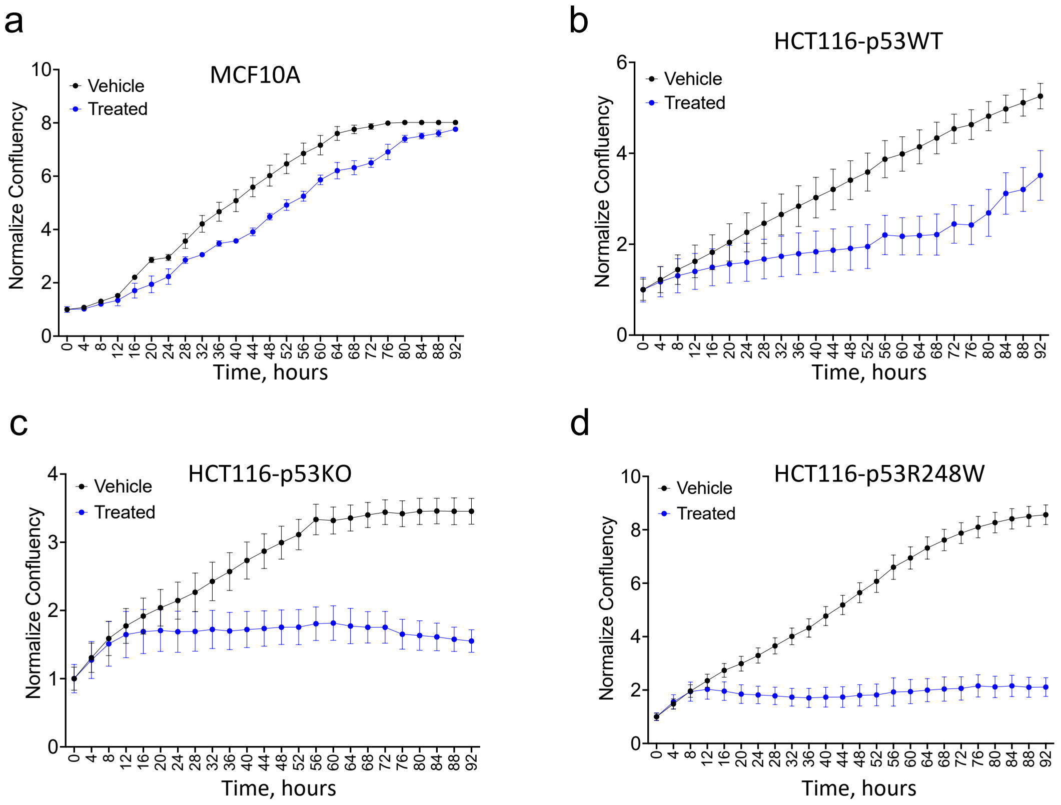


**Suppl. Fig. 6. Recovery of cell growth after TAS102-talazoparib treatment.** (**a-c**) Cell growth curves of MCF10A, HCT116-p53WT, HCT116-p53KO, and HCT116-p53R248W/- cell lines treated with 500nM of TAS102 plus 100nM of talazoparib for 24hrs, followed by drug washout, and monitored for cell growth over time using live imaging system. All experiments were repeated at least 2 times.

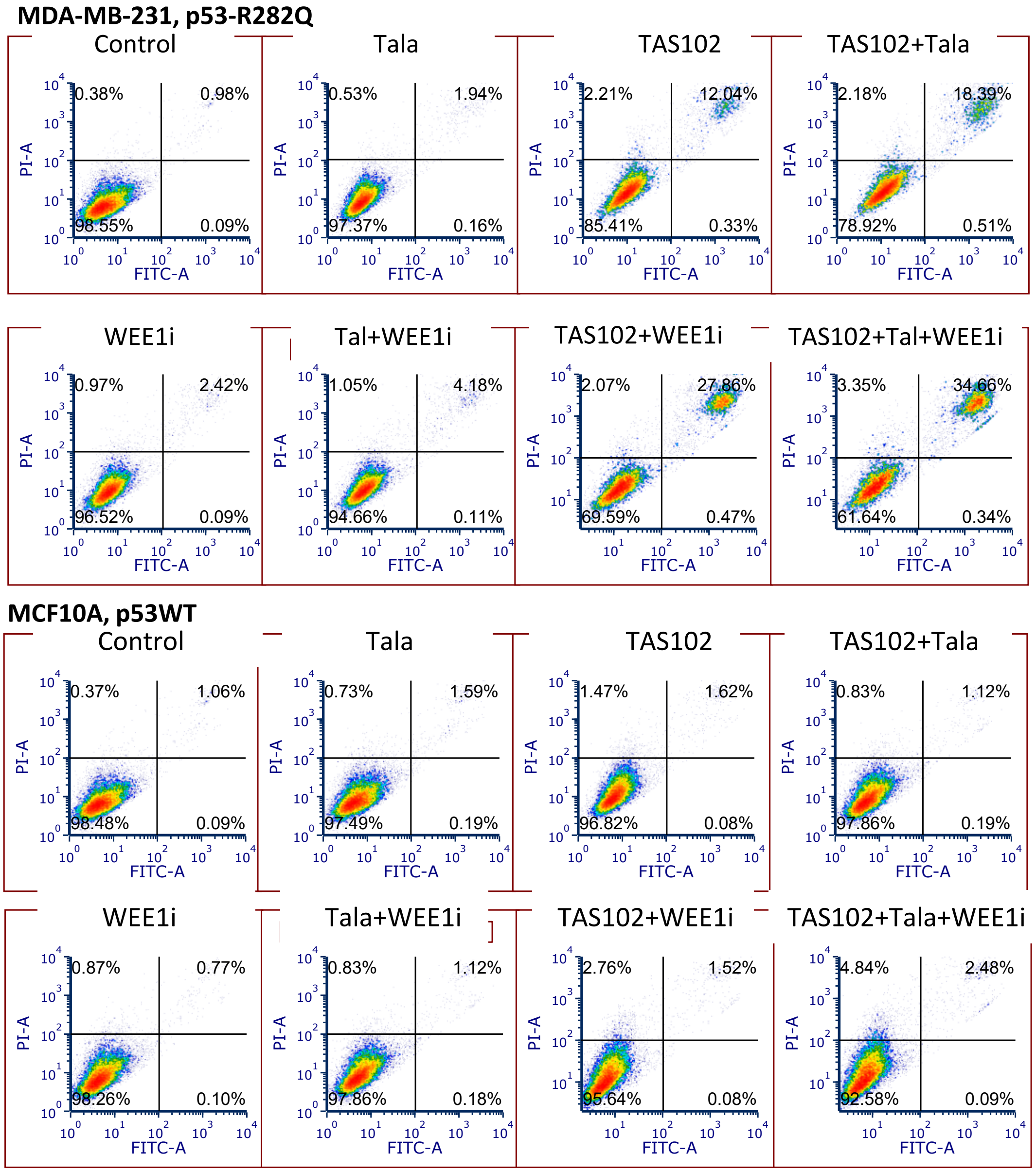


**Suppl. Fig. 7. Cell viability growth after TAS102-talazoparib treatment.** Cell viability was evaluated by flow cytometry analysis of cells stained with propidium iodide (PI) and annexin-V. Cells were treated with vehicle-control or 500nM TAS102 plus 100nM talazoparib for 24hours, then 24-hr washout that followed by incubation with 250nM WEE1i for 48hours, where it is indicated. All experiments were repeated at least 2 times.

a

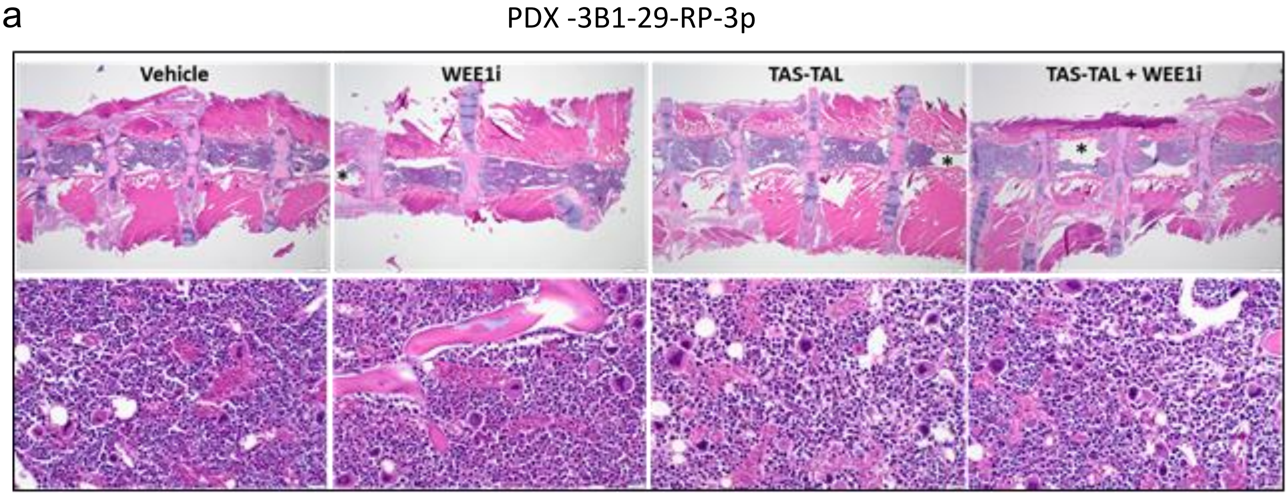


b

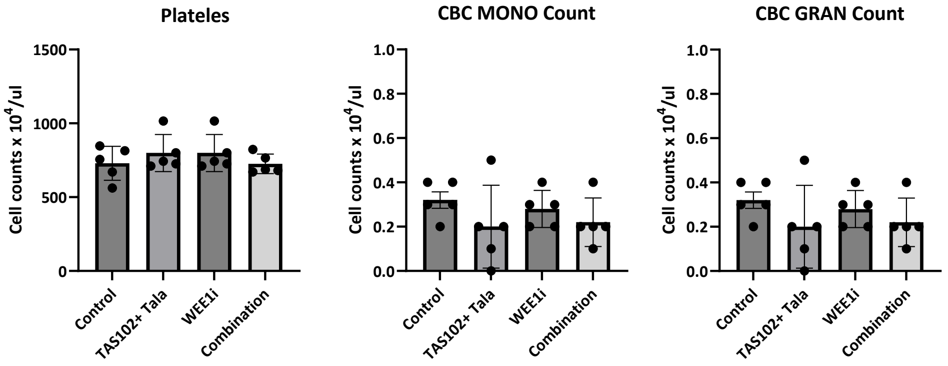


**Suppl. Fig. 8. Effects of treatments on bone marrow and peripheral blood.** (**a**) Sternal bone marrow was assessed for histology using H&E stain in three mice per each group, representative images are shown. Areas of apparent marrow dropout (marked with an asterisk) were attributed to slide preparation artifacts. Mice were treated as described for PDX -3B1-29-RP-3p in Figure 7. (**b**) Peripheral blood was evaluated in five mice per each group. No statistically significant differences were noted. Comparisons were performed using one-way ANOVA, n=5.
